# Biochemical characterization of the *Escherichia coli* surfaceome: A focus on type I fimbriae and flagella

**DOI:** 10.1101/2024.01.14.575581

**Authors:** DW Kavanaugh, A Sivignon, Y Rossez, Z Chouit, C Chambon, L Beal, M Hébraud, Y Guerardel, H Nguyen, N Barnich

## Abstract

The *Escherichia coli* surfaceome consists mainly of the large surface organelles expressed by the organism to navigate and interact with the surrounding environment. The current study focuses on type I fimbriae and flagella. These large polymeric surface organelles are composed of hundreds to thousands of subunits, with their large size often preventing them from being studied in their native form. Recent studies are accumulating which demonstrate the glycosylation of surface proteins or virulence factors in pathogens, including *E. coli*.

Using biochemical and glycobiological techniques, including biotin-hydrazide labelling of glycans and chemical and glycosidase treatments, we demonstrate i) the presence of a well-defined and chemically resistant FimA oligomer in several strains of pathogenic and non-pathogenic *E. coli*, ii) the major subunit of type I fimbriae, FimA, in pathogenic and laboratory strains is recognized by concanavalin A, iii) standard methods to remove *N*-glycans (PNGase F) or a broad-specificity mannosidase fail to remove the glycan structure, despite the treatments resulting in altered migration in SDS-PAGE, iv) PNGase F treatment results in a novel 32 kDa band recognized by anti-FliC antiserum.

While the exact identity of the glycan(s) and their site of attachment currently elude detection by conventional glycomics/glycoproteomics, the current findings highlight a potential additional layer of complexity of the surface (glyco)proteome of the commensal or adhesive and invasive *E. coli* strains studied.

## Introduction

*Escherichia coli* (*E. coli*) uses a range of virulence factors to promote their adhesion, colonization, and invasion of human eukaryotic epithelial cells as well as immune cells^1^. To facilitate adhesion to sites, such as the small intestine or urethra, pathogenic *E. coli* use adhesins most commonly in the form of type I fimbriae (also referred to as pili), among others^2^. Type I fimbriae have been extensively studied during the last 70 years^3–6^.

They are composed of 500 to 3,000 FimA subunits, and are generally thought to be terminated by FimF, FimG, and FimH, the latter allowing interaction with host and immune cell structures primarily via the mannose glycan^5,7^. Subunit assembly is facilitated by donor strand complementation, with an extension of each subunit non-covalently inserted into a pocket of the preceding subunit in the assembly. While FimF, FimG, and FimH subunits are easily dissociated from the polymer via SDS treatment, FimA resists SDS treatment, chaotropic salts including guanidium chloride and urea, and trypsin^3^. Importantly, the terminal localization of FimF, FimG, and FimH has been questioned, with limited studies giving evidence for the potential distribution of the minor subunits along the pili shaft, creating potential weak points. Abraham *et al*. provided evidence for this phenomenon using confocal microscopy and immunogold labeling, demonstrating the presence of FimH along the fimbrial shaft, while Ponniah *et al*. demonstrated that freeze-thaw-induced pilus fragmentation leads to the emergence of high-affinity mannose-binding groups, proposed to be the exposure of non-terminal FimH^8^. FimA subunits are readily dissociated following mild acid hydrolysis (> pH 2, 99°C, 5 min), enhancing protein detection and, in specific cases, enabling the detection of reducing sugars^3^. However, as type I fimbriae are large biopolymers, their study is intrinsically complicated, and little progress has been made in studying their intact structure in recent years.^9^

Despite their widespread presence and conserved sequence homology between pathogenic and non-pathogenic *E. coli* strains, both fimbriae and flagella may contribute to strain differentiation^10–12^, further reinforcing the interest for vaccine development. For example, with respect to Crohn’s disease-associated strains, defined as adhesive and invasive *E. coli* (AIEC), point mutations in the type I pili adhesive subunit, FimH, can be used to further categorize strains based on their increased ability to adhere to an intestinal ligand carrying the cognate sugar (mannose) of the pili adhesin lectin^13^. Additionally, Duncan *et al*. have demonstrated that FimH-binding to mannosylated structures is modulated by the host origin of the FimA shaft^12^ in which expression of a *Klebsiella* FimH on a pili shaft of *E. coli* origin altered the binding selectivity compared with that of an *E. coli* fimbriae, thus highlighting the important contribution of the fimbrial shaft and not only the terminal lectin.

Until recently, little attention has been given to the possibility of post-translational modification of the fimbrial subunits and its potential impact. Originally, a summary of research on type I fimbriae carried out between 1951-1965 by Brinton stated that chemical analysis of type I fimbriae revealed no detectable amounts of carbohydrate^6^. More recently, however, two studies have highlighted the potential glycosylation of pili - McMichael and Ou^3^, and a patent granted in 2002^11^. Later, another study identified FimH as a candidate glycoprotein interacting with the GlcNAc-binding lectin, WGA^14^. Recent findings indicate that glycan modifications of surface proteins may be associated with altered motility as well as interactions between bacteria and with host epithelial and immune cells^15–18^.

Forty years have passed since McMichael and Ou demonstrated the detection of reducing sugars present on purified fimbriae only after acid hydrolysis, with their additional findings proposing the presence of *O*-glycans and potentially other uncharacterized glycans^3^. The identity of *E. coli* fimbriae-associated glycans has been hypothesized based on the work of Palacios-Palaez *et al*.^11^ using lectin profiling, albeit limited to uropathogenic *E. coli* strains. The latter work implies the presence of mannose, *N*-acetylglucosamine, and sialic acid, but has not been further reported.

Despite indications of glycan post-translational modifications of fimbriae, *E. coli* lack the well-defined and characterized glycosylation systems currently identified among Gram-negative organisms, including *Campylobacter, Neisseria*, and others^19–23^. However, recent studies show promise in identifying *E. coli* glycosylation enzymes, with those of the LPS biosynthesis pathway playing a dual role. In particular, Benz and Schmidt demonstrated the modification of the adhesin responsible for diffuse adhesion (AIDA-I) with an average of 19 heptose molecules obtained from the pathway of LPS synthesis by AAH (autotransporter adhesin heptosyltransferase)^16^. More recently, Thorsing *et al*.^24^ characterized the heptosylation of the ETEC H10407 protein, YghJ, with the non-glycosylated version of the protein expressed in the *E. coli* K-12 MG1655 strain lacking the heptosyltransferase gene, *hldE*, further underlining the dual nature of the LPS synthesis pathways as a source of both enzymes and substrate for glycosylation of *E. coli* proteins.

This study examines the large surface organelles of several bacterial strains in both their native and depolymerized states. Given the complex and challenging nature of studying type I fimbriae, this study seeks to explore their potential for post-translational glycan modifications and, in doing so, has identified novel properties and features associated with type I fimbriae and flagella of *E. coli*. The main results of this study highlight the existence of a predominant FimA-containing pili unit of around 90 kDa, in several pathogenic (AIEC) and non-pathogenic strains (*E. coli* K-12), which is resistant to SDS-mediated dissociation, and confirm the presence of FimA-associated glycosylation, which is resistant to several chemical and enzymatic treatments.

## Results

### Surface protein purification and general characterization

The major surface proteins of the panel of *E. coli* strains were initially extracted by mechanical shearing, as previously described^9^. However, to limit the impact of mechanical stress on fimbria and flagella polymers, extraction was performed as per Barnich *et al*., via thermal treatment^25^, demonstrating no significant effects on yields of extracted surface proteins (not shown).

Total surface protein extracts of a panel of strains were profiled by Ponceau S (Fig. 1A) and Alcian blue staining (Fig. 1B). Ponceau S staining demonstrates the various profiles of extracted surface proteins, while Alcian blue staining demonstrates the presence of acidic polysaccharides in the extracted samples appearing as well-defined bands. This is in contrast to smearing which is associated with migration of capsular polysaccharides. Of note, Alcian blue staining resulted in negative staining of the non-stained proteins. The majority of adherent and invasive strains stained for Alcian blue at ∼72 kDa, with AIEC CEA218U (Fig. 1B, lane 12) demonstrating the most intense and variable staining of these AIEC strains. The two AIEC strains not staining at 72 kDa also demonstrate a lack of staining at this size in the Ponceau S image. Additionally, POP18-M0 (Fig. 1B, lane 8) stains at 36 kDa, and POP179-M0 (Fig. 1B, lane 11) at 38 kDa. *E. coli* MG1655 and *E. coli* C600 stain at ∼55 kDa, while *E. coli* HS (Fig. 1B, lane 4) stains at ∼38kDa, with Alcian blue staining generally coinciding with the strongest bands in the Ponceau S staining.

**Figure 1.**
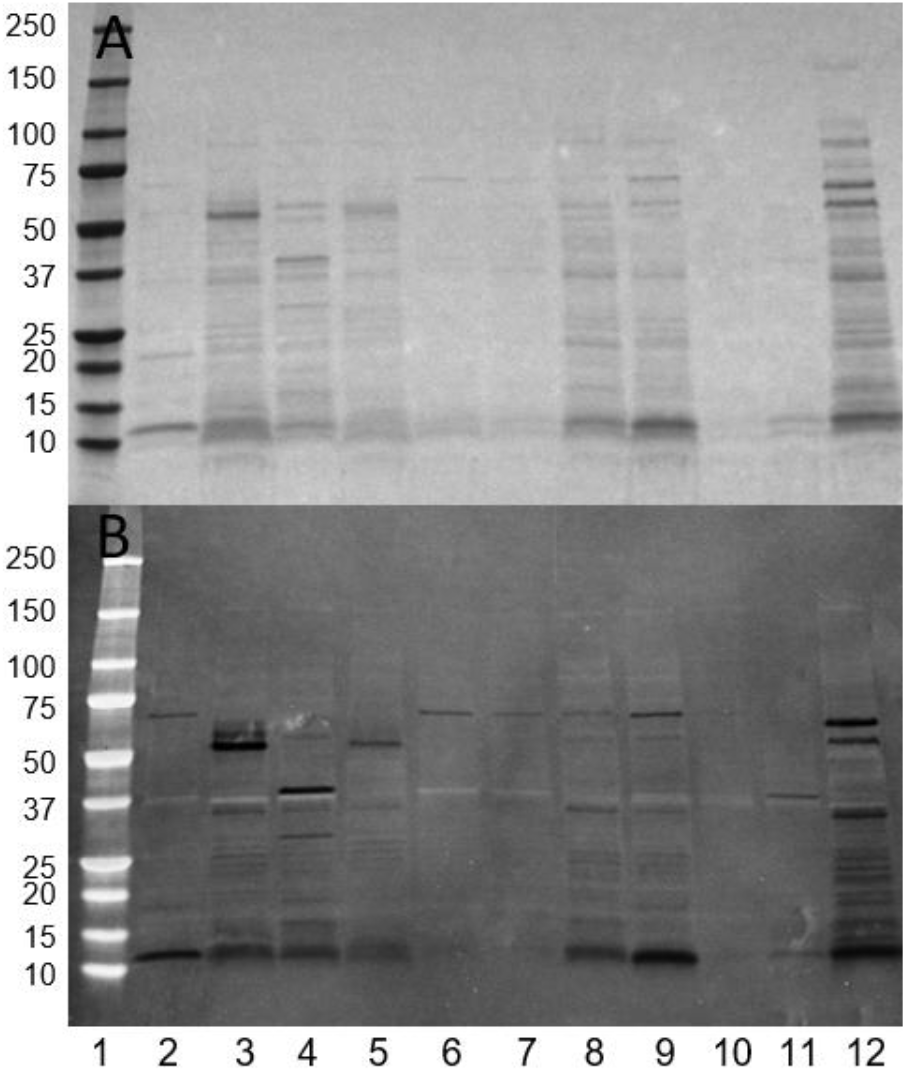
Visualization of released surface proteins of a panel of *E. coli* strains. (A) Ponceau S. (B) Alcian blue staining for acidic polysaccharides (dark bands). Lanes 1. Molecular weight marker, 2. AIEC LF82, 3. MG1655, 4. *E. coli* HS, 5. *E. coli* C600, 6. AIEC LF82 Δ*fimA*, 7. AIEC LF82 Δ*fimH*, 8. POP18-M0, 9. POP90-M0, 10. POP135-M0, 11. POP179-M0, 12. CEA218U.

To limit the potentially deleterious effects of TCA on the released proteins and to aid in enrichment of type I fimbriae and flagella, as well as other present surface proteins (including OmpA and OmpC), released *E. coli* surface proteins were precipitated by 30% ammonium sulfate, collected, and resuspended in milli-Q water. The samples often remained cloudy and resistant to complete re-solubilization. In both, clarified and turbid samples, fimbria/FimA and FliC represent the primary proteins detected by Coomassie blue staining. However, removal of the insoluble material prior to Western blot analysis results in a significant decrease in detection of pili polymer fragments or acid-hydrolysis-released FimA subunits (Fig. 2; lane 2 (non-clarified) versus lane 4 (centrifuged)) as well as a reduction in GNA staining. As the goal was to investigate the potential differences in polymer configurations or the presence of modifications, it was important to conserve the fimbrial fragments, avoiding removal of potentially important molecules by over-purification.

**Figure 2.**
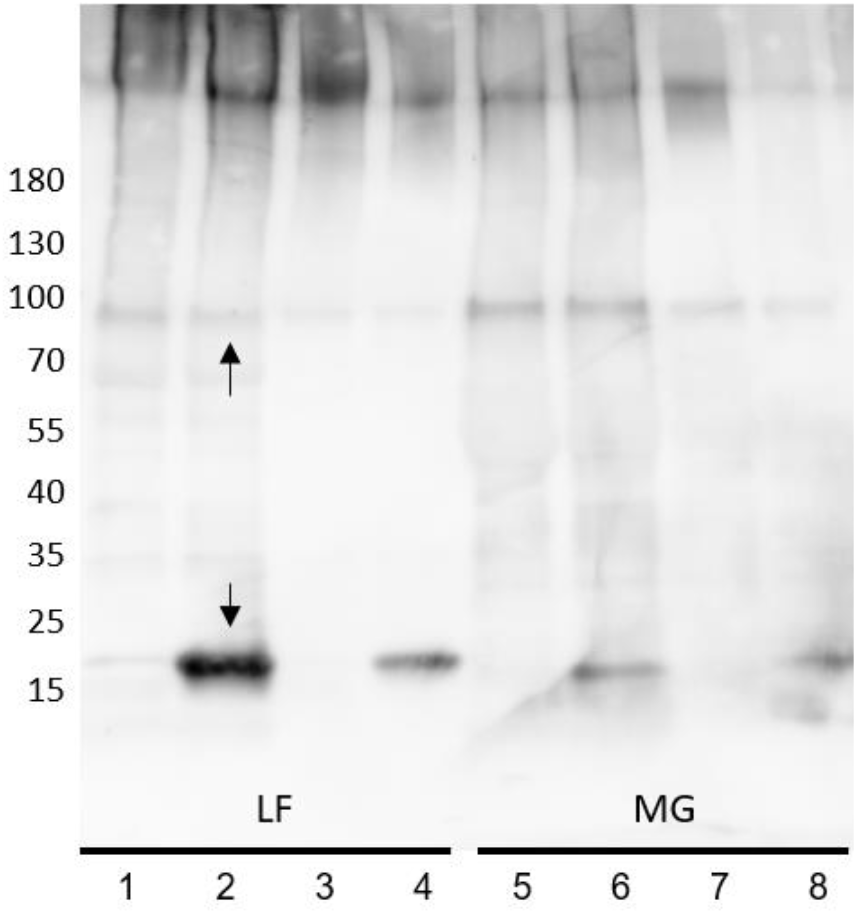
Anti-type I fimbria staining of released *E. coli* surface proteins. Lane 1, 2, 5, 6: Crude extracts; Lane 3, 4, 7, 8: Clarified supernatants. Samples non-acid-hydrolyzed: lanes 1, 3, 5, 7. Acid-hydrolyzed samples: lanes 2, 4, 6, 8. Upper arrow denotes the 90 kDa FimA oligomer, the lower arrow denotes the FimA subunit.

### Narrowing the focus to type I fimbriae

Use of MgCl_2_ to further enrich the surface protein extracts for type I fimbria and flagella was carried out. Initially, the study of type I fimbria included analysis of intact fimbria employing typical SDS-PAGE conditions, and depolymerized FimA subunits by means of mild acid hydrolysis. Western blot of surface extracted proteins with anti-type I fimbria antiserum revealed the expected intense staining of FimA monomers at 18 kDa (Figure 2, lower arrow) for AIEC LF82 following mild acid hydrolysis. Interestingly, under non-hydrolyzed conditions, a distinct band of approximately ∼90 kDa stains for anti-type I fimbriae (Figure 2, upper arrow) indicating the presence of a conserved protein oligomer which is resistant to depolymerization by heat and SDS (Fig. 2, Fig. 3 – asterisk*). Accordingly, incubation with anti-FimF, anti-FimG, or anti-FimH did not stain the 90 kDa band (data not shown), confirming the expected absence of these fimbrial subunits in the high molecular weight polymers following preparation in Laemmli buffer containing SDS (as previously established by Jones et al., ^26^). As such, the stained band likely represents a FimA homopentamer, corresponding to 5 × 18 kDa. This major band represents the minimal FimA oligomer unit to be resolved under SDS-PAGE conditions in these sample preparations, implying a minimum span of 5 FimA subunits separated by SDS/beta-mercaptoethanol-sensitive subunits. Given the absence of detection of a FimA subunit at 18 kDa, it is hypothesized that these stretches of FimA are likely separated by complexes of FimF, FimG, FimH.

**Figure 3.**
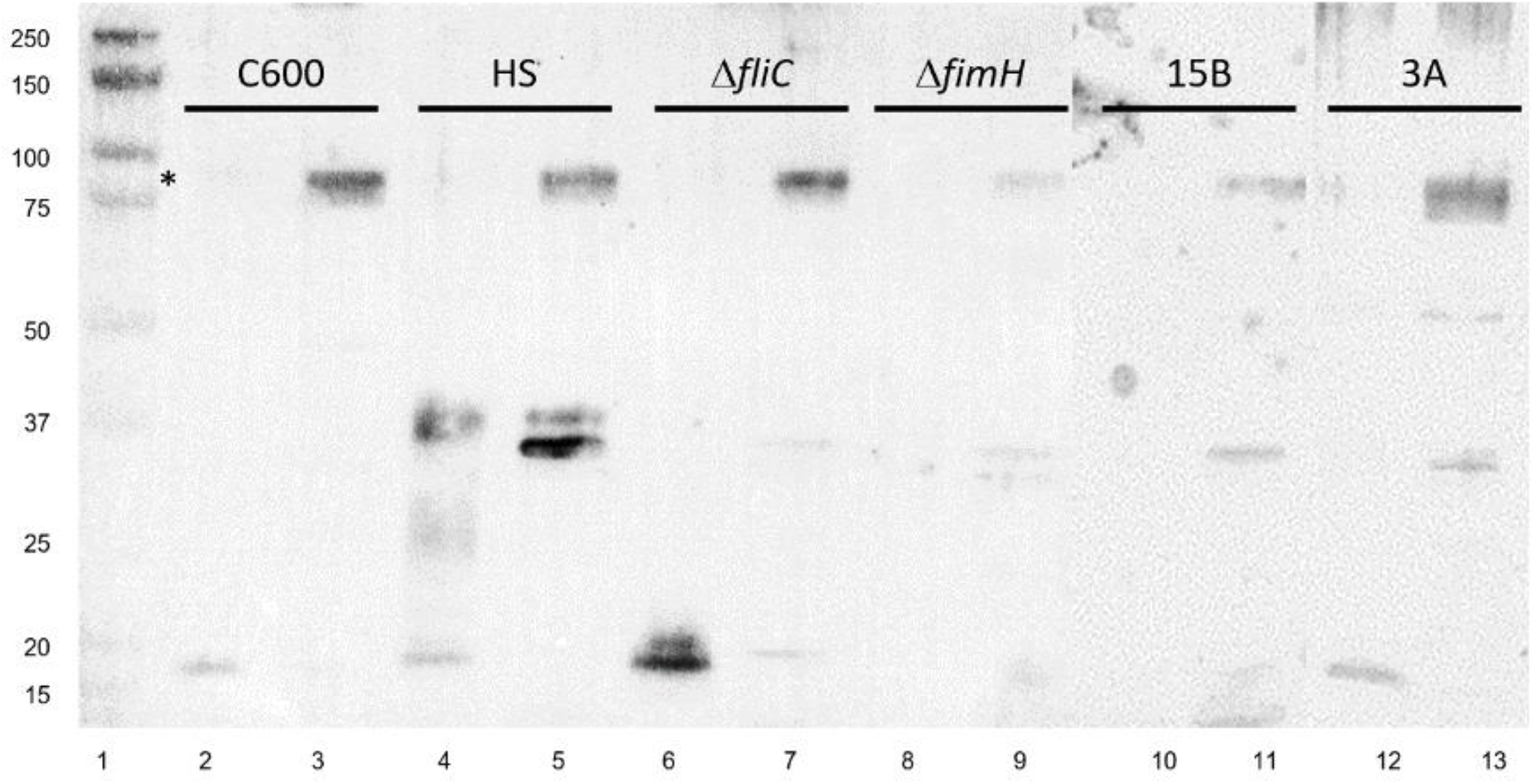
Anti-type I fimbria staining of panel of *E. coli* strains. Lane 1: Molecular weight ladder; lanes 2,3: *E. coli* C600; lanes 4,5: *E. coli* HS; lanes 6,7: LF82Δ*fliC*; lanes 8,9: LF82 Δ*fimH*; lanes 10,11: AIEC CEA614S (15B); lanes 12,13: AIEC CEA218U (3A). Splice of membranes treated identically. * FimA oligomer present under non-hydrolyzed conditions. Strains are shown in pairs as acid-hydrolyzed and non-hydrolyzed, respectively.

The 90 kDa band is present across all other strains tested from our collection following enrichment by MgCl_2_ treatment, including *E. coli* C600 and HS, and other strains of AIEC, including CEA218U and CEA614S (Fig. 3), thus does not appear to be strain specific given the anti-type I fimbria serum used for staining.

### Depolymerization of type I fimbria via chemical treatment

To facilitate purification and, later, interaction analysis of FimA monomers, two approaches were tested for the release of FimA subunits. First, reduction and alkylation were tested via incubation with dithiothreitol (DTT) and iodoacetamide (IAA), and secondly, incubation in saturated guanidine hydrochloride (GuHCl). These approaches were employed to avoid the potentially degradative effects of acid hydrolysis which is typically employed for FimA subunit release.

FimA possesses a cysteine bridge between residues 46-86. To open the tightly coiled fimbriae structure, reduction and alkylation of samples was carried out. As seen in Fig. 4A, in both strains, LF82 and MG1655, the treatment enabled the detection of the FimA monomer (18 kDa and 15 kDa, respectively) by Coomassie blue staining, without the prior need of acid hydrolysis. In the case of GuHCl treatment of fimbriaenriched surface protein extracts, the release of FimA subunits is much more efficient than by reduction and alkylation alone, as determined by anti-type I fimbriae detection (Fig. 4B), yet produces FimA tetramers to a lesser extent (Fig. 5). Additionally, reduction and alkylation results in the presence of FimA dimers, as revealed by Western blot. GuHCl treatment leaves the cysteine bridge intact, thus migrating further as a more compact structure, while reduction and alkylation breaks the disulfide bridge resulting in an opened structure, migrating at a larger perceived size (Fig. 4B). To further probe the effects of reducing agents on the structural composition of FimA oligomers, we tested GuHCl-treated samples in the presence or absence of both reduction/alkylation and betamercaptoethanol (B-ME) independently or together. Of note, the loading buffer contains SDS, which has been shown to release the FimF, FimG, and FimH subunits from the full-length fimbriae polymer. Treatment with B-ME alone results in no change to FimA tetramers (∼72 kDa), while FimA monomers are present due to the initial GuHCl treatment (Fig. 5, column 2: absence of B-ME and DTT/IAA versus column 3: presence of B-ME and absence of DTT/IAA). As mentioned above, reduction and alkylation results in FimA dimers and monomers with the disappearance of the FimA tetramers, implying the conversion of FimA tetramers to FimA dimers. Finally, addition of B-ME to reduced and alkylated samples results in only FimA monomers, disrupting the association between FimA dimers (Fig. 5). Notably, addition of DTT to Laemmli buffer containing beta-mercaptoethanol is not sufficient for complete dissociation (data not shown), indicating the requirement of either GuHCl treatment or alkylation, or both to achieve dissociation of tetramers and dimers.

**Figure 4.**
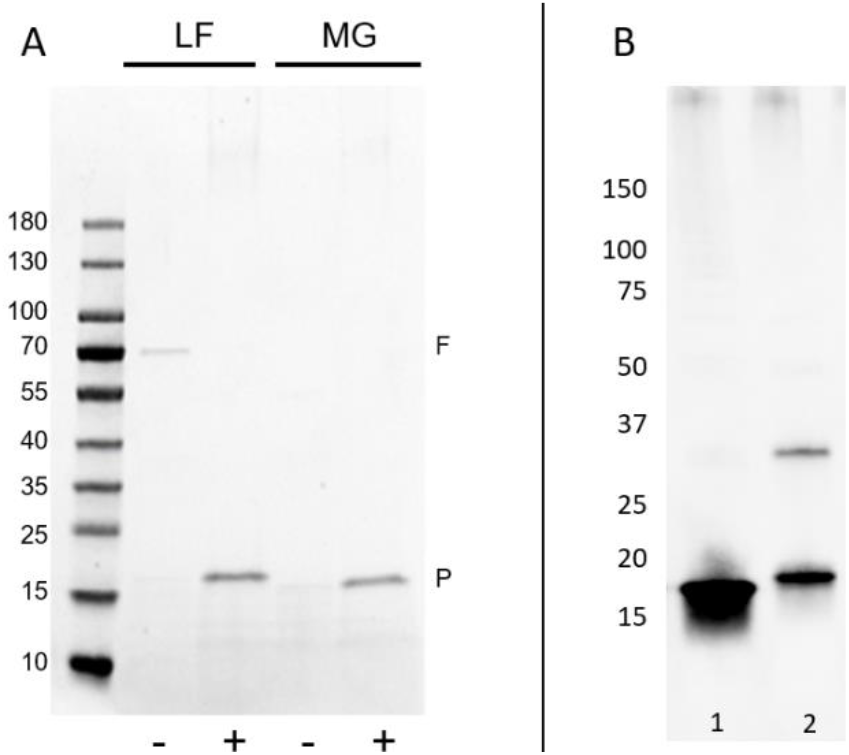
Influence of reduction and alkylation or guanidine hydrochloride treatment on FimA release and detection. (A) Coomassie blue staining of reduced and alkylated (+) or non-treated (-) fimbria-enriched surface protein extracts. Samples were not acid-hydrolyzed to aid the release and staining of FimA subunits. F = flagella (FliC), P = pili (fimbria; FimA), LF – AIEC LF82, MG – *E. coli* K-12 MG1655. (B) Guanidine HCl treatment versus reduction and alkylation of type I fimbria-enriched protein extracts from LF82. Type I fimbriae detection by anti-type I fimbria antiserum. Lane 1: GuHCl-treated, lane 2: reduced and alkylated.

**Figure 5.**
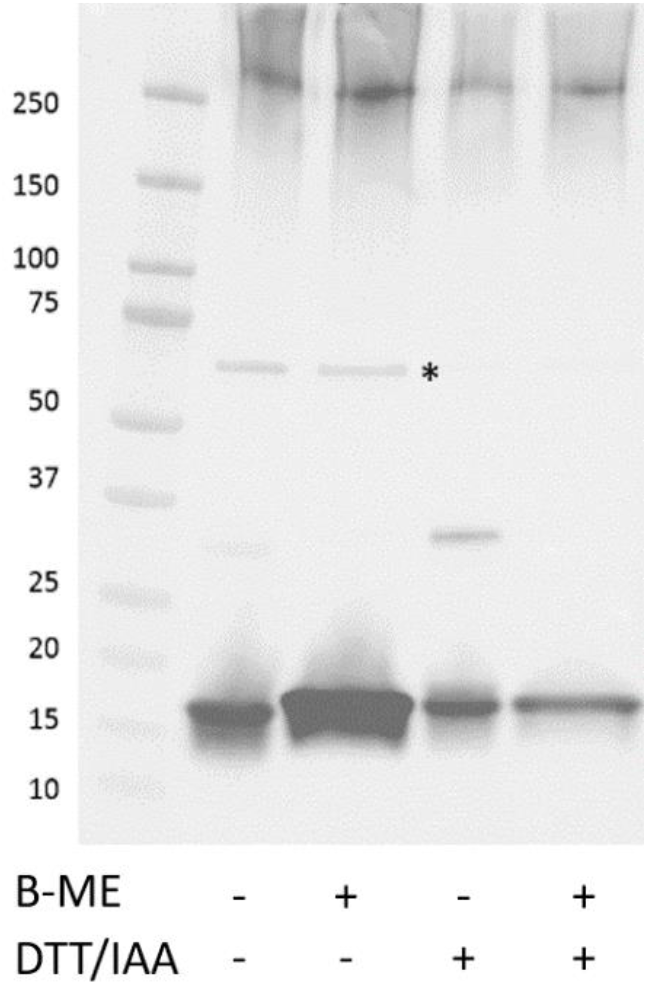
Effect of reducing agents on the depolymerization of FimA oligomers. GuHCl-treated LF82 surface extracted proteins in the absence or presence of varying reducing agents were resolved by SDS-PAGE followed by detection of type I fimbria. * denotes FimA tetramer. B-ME – Beta-mercaptoethanol, DTT/IAA – reduction and alkylation.

Given the various steps involved in depolymerization of type I fimbria and the eventual demonstration of the interaction between FimA and the lectin, concanavalin A (ConA), we propose a graphical summary of the evidence presented (Fig. 6).

**Figure 6.**
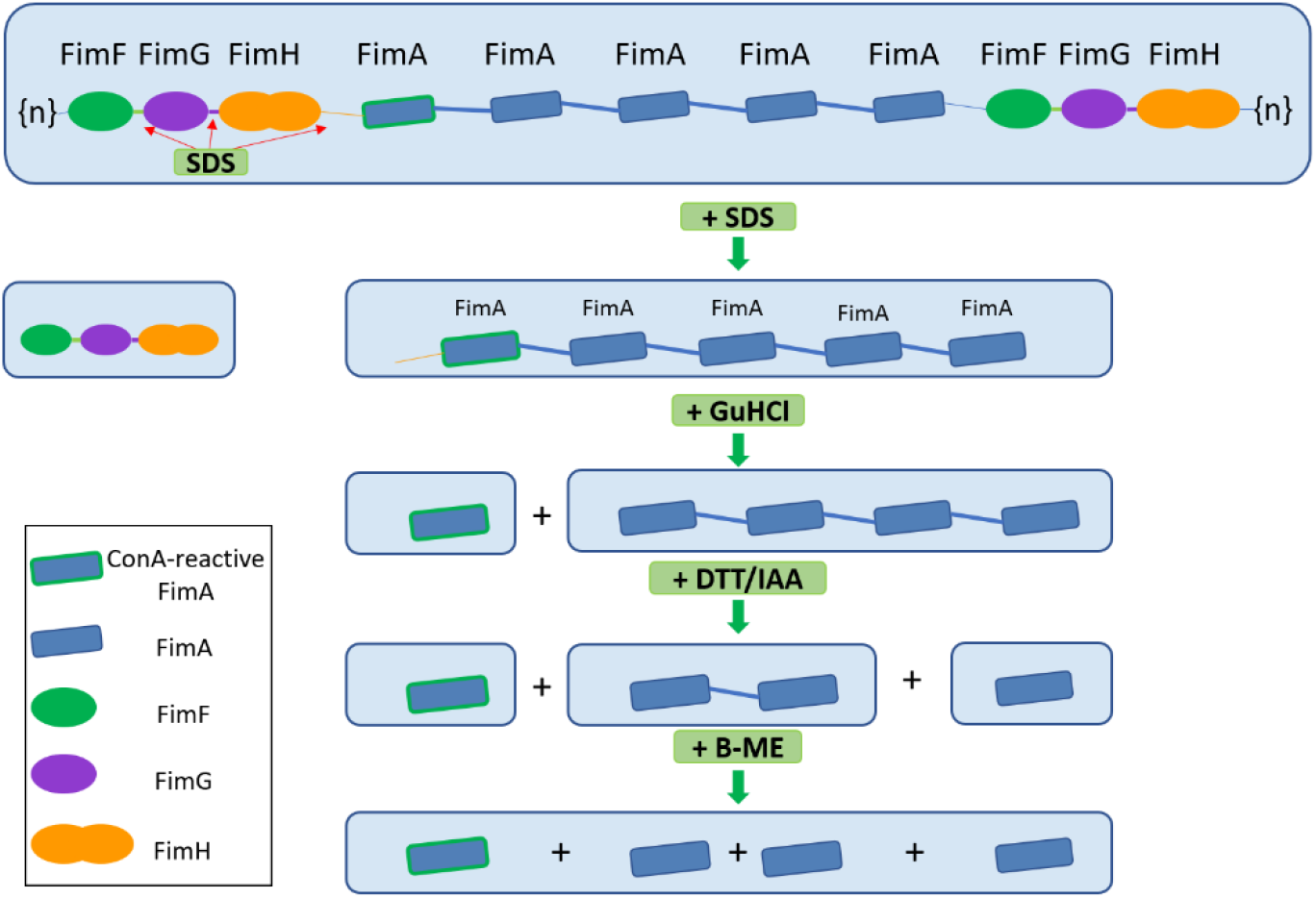
Graphical scheme of the depolymerization of type I fimbria following several stepwise chemical treatments. FimF, G, and H are removed by SDS and heat. GuHCl treatment further releases ConA-reactive FimA subunits. Reduction and alkylation (DTT/IAA) produces FimA dimers and monomers. Lastly, beta-mercaptoethanol dissociates the remaining FimA dimers into monomers. Beta-mercaptoethanol is able to release FimA subunits, however, the origin of these subunits does not appear to be the FimA tetramers, but likely the high molecular weight polymer migrating above 180 kDa.

### Lectin staining

#### Concanavalin A (ConA

*E. coli* surface protein extracts, either acid hydrolyzed (AH) to release FimA subunits or left untreated (NT) to analyze intact fimbriae, were stained with either ConA, GNA, or WGA. ConA strongly stains the released FimA subunits, but does not stain samples that were not acid-hydrolyzed (Fig. 7C) implying the presence of a glycan embedded within the quaternary structure of the fimbriae. ConA and anti-type I fimbria staining overlap identically during the detection of released FimA subunits (Fig. 7B + C). However, as ConA is reported as recognizing glucose, mannose, or L/D-glycero-manno-heptoses^27^, the identity of the attached glycan has not yet been confirmed. Sugar inhibition assays were carried out with glucose and mannose in an attempt to block ConA binding to FimA, however, this was unsuccessful, indicating the presence of an alternative glycan (results not shown). As further confirmation of the ConA staining of FimA subunits, lectin blotting of the AIEC LF82 wild-type, along with isogenic mutants for the *fimA* and *fliC* genes was conducted, demonstrating an absence of ConA staining when the strain no longer expresses *fimA*, but not when *fliC* is deleted (Fig. 8).

**Figure 7.**
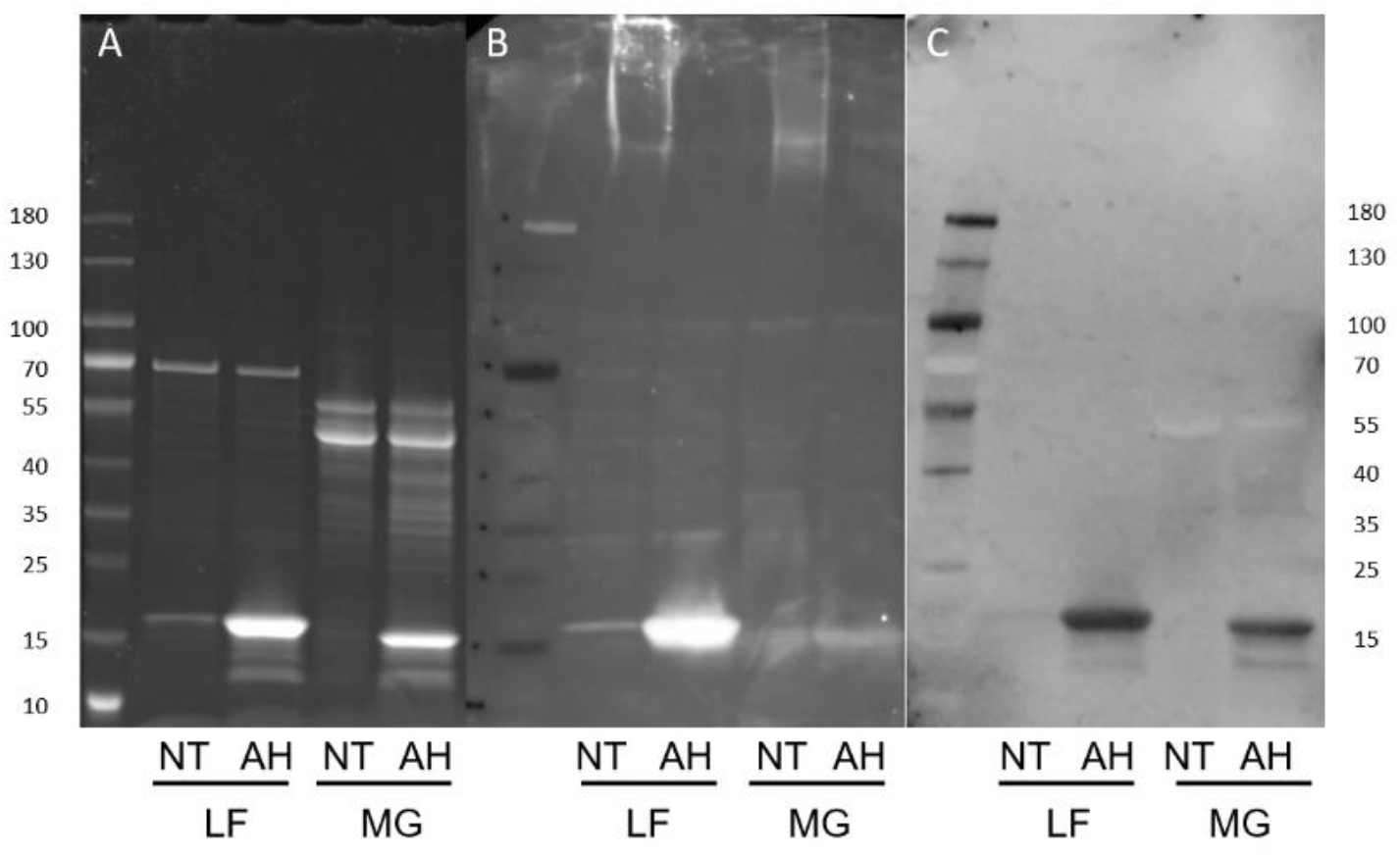
Protein and lectin staining of released *E. coli* surface proteins. NT: Non-treated, AH: Acid-hydrolyzed, LF: AIEC LF82. MG: *E. coli* MG1655. (A) Coomassie blue stain, (B) Anti-type I fimbriae-FITC staining, (C) ConA-HRP staining.

**Figure 8.**
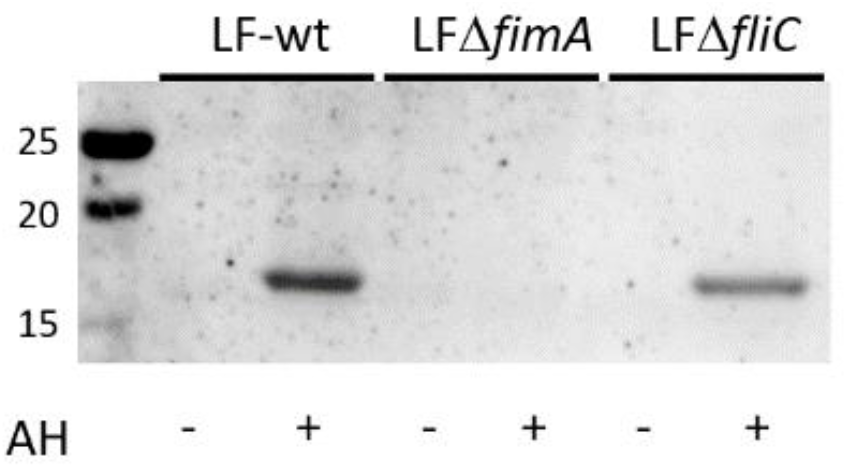
ConA staining of type I fimbriae-enriched LF82 surface protein extracts. ConA staining is negative in the absence of type I fimbria. AH – acid hydrolysis treatment.

Comparing the standard acid hydrolysis treatment against GuHCl treatment, acid hydrolysis appears to be less effective in releasing ConA-reactive FimA subunits (Fig. 9), even when the GuHCl samples were prepared in the absence of beta-mercaptoethanol (B-ME). Additionally, acid hydrolysis produces a FimA monomer of the same size as that produced following reduction and alkylation, indicating that the treatment also disrupts the disulfide bridge of the monomer.

**Figure 9.**
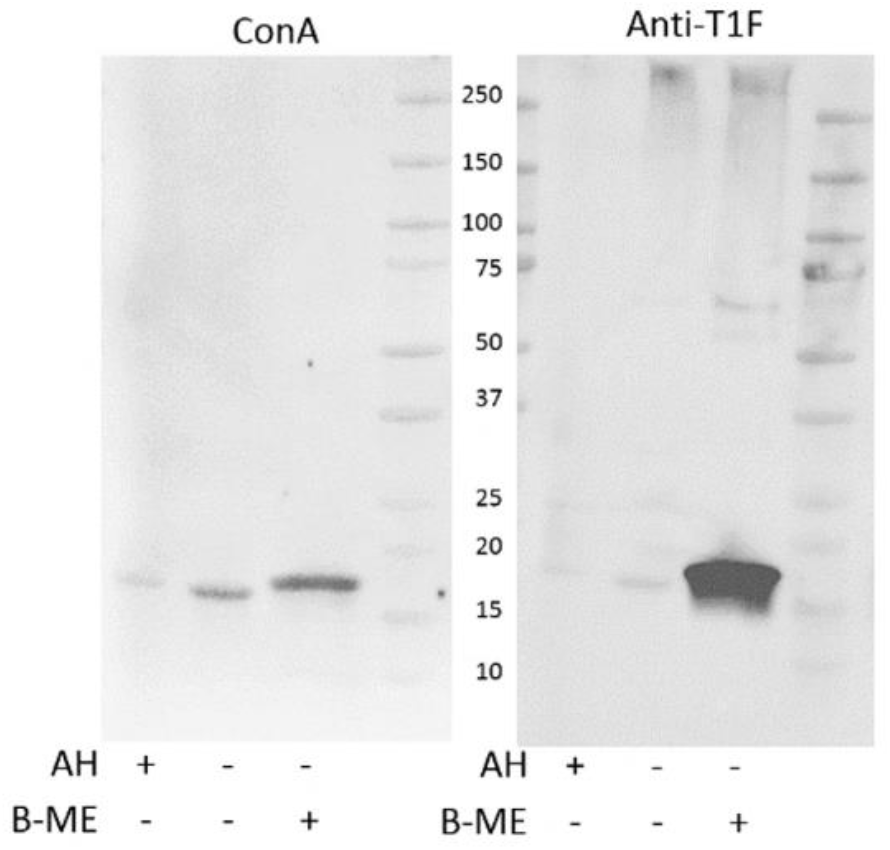
Guanidine HCl-treatment efficiently releases ConA-reactive FimA subunits. Extracted surface proteins of LF82 were treated with GuHCl and analyzed by SDS-PAGE and Western blot. Acid *hydrolyzed* (AH) sample was included as positive control for FimA subunits. GuHCl-treated samples were prepared in the presence or absence of beta-mercaptoethanol (B-ME). The blot was probed with ConA (A), followed by anti-type I fimbria antiserum (Anti-T1F) (B).

The addition of B-ME does not significantly increase the amount of ConA-reactive FimA, however, it greatly increases staining by anti-type I fimbria detection (Fig. 9, ConA staining (left image) versus anti-T1F staining (right image). This leads to the hypothesis that: 1) reduction of the disulfide bond enables better antibody access, or 2) there exist at least two forms of FimA subunits – ConA-reactive, released by GuHCl, and those not recognized by ConA and released in the presence of B-ME. To confirm the efficacy of GuHCl treatment to release ConA-reactive FimA monomers, untreated and GuHCl-treated proteins were dot-blotted to a nitrocellulose membrane in the absence of beta-mercaptoethanol, DTT, and SDS and probed with ConA. The GuHCl-treated sample demonstrated ConA binding, while the untreated sample did not, despite equal loading of protein determined by Ponceau S staining (data not shown). Together, these results indicate a differential response of FimA oligomers in the presence of different reducing agents or conditions.

### GNA and WGA staining

Galanthus nivalis (GNA; mannose) lectin staining was comparatively weak, requiring overnight lectin incubation and Clarity max ECL reagent for visualization, but consistently present in the region between the wells of the gel and ∼75 kDa gel marker, running as a smear, and overlapping with the anti-type I fimbria staining of the poorly resolved non-hydrolyzed fimbrial polymer (Fig. 10A). Wheat germ agglutinin (WGA; sialic acid and N-acetylglucosamine) staining was observed strongly in the MG1655 surface protein extract (Fig. 10B), interacting with a lower molecular weight structure of between 12-14 kDa. This structure has yet to be identified, as antibody staining for FimA, FimF, or FimG were not superimposable with this staining.

**Figure 10.**
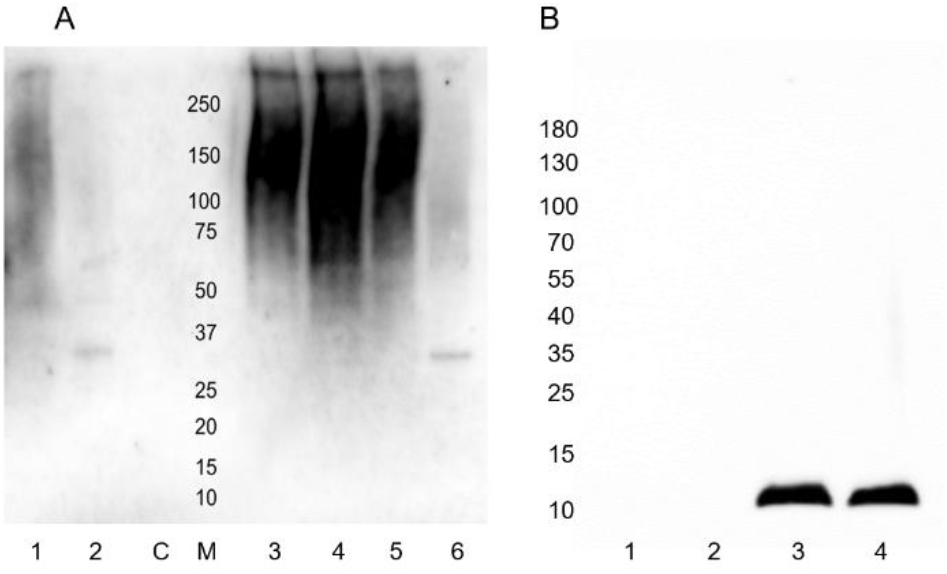
Lectin staining of released *E. coli* surface proteins. (A) GNA-HRP staining. Lane 1: Mock-treated LF82; Lane 2: PNGase F-treated LF82; Lane 3: Acid hydrolyzed *E. coli* MG1655; Lane 4: Untreated MG1655; Lane 5: Mock-treated MG1655; Lane 6: PNGase F-treated MG1655. C: PNGase F enzyme only control; M: Molecular weight ladder. (B) WGA-biotin staining. Lane 1: LF82 non-hydrolyzed; Lane 2: LF82 acid-hydrolyzed; Lane 3: MG1655 non-hydrolyzed; Lane 4: MG1655 acid-hydrolyzed.

### Lectin-independent glycan detection

Lastly, to confirm the presence of glycosylated structures in the fimbria-enriched surface protein extracts, attached glycans were labelled with biotin-hydrazide, followed by detection with streptavidin-horseradish peroxidase/ECL detection. This staining of the resolved proteins on the PVDF blot demonstrates the presence of a glycan structure on the FimA subunit, when either acid hydrolyzed or depolymerized via GuHCl treatment, while the untreated sample does not demonstrate staining (Fig. 11B). Additionally, negative controls, including soybean trypsin inhibitor, as well as the molecular weight ladder, were absent of staining. Interestingly, in the *E. coli* K-12 MG1655 strain fimbria-enriched surface protein extract, there is staining of the appropriate size for the FimA subunit (15 kDa, predominantly in the GuHCl-treated sample), but interestingly also staining of multiple bands in the approximate range of 36-80 kDa, in all sample treatments, including untreated and acid hydrolyzed, consistent with sizes above and below the predominant Ponceau S staining for the FliC protein. These results are in accordance with the ConA staining for AIEC LF82 FimA, wherein staining only occurs following either acid hydrolysis or GuHCl treatment, pointing to the presence of a sterically protected glycan. Regarding MG1655, the biotin-hydrazide staining confirms the ConA staining for the FimA subunit, as for LF82, but also lends evidence to the presence of additional distinct glycoproteins or a glycoprotein oligomer in the surface protein preparation.

**Figure 11.**
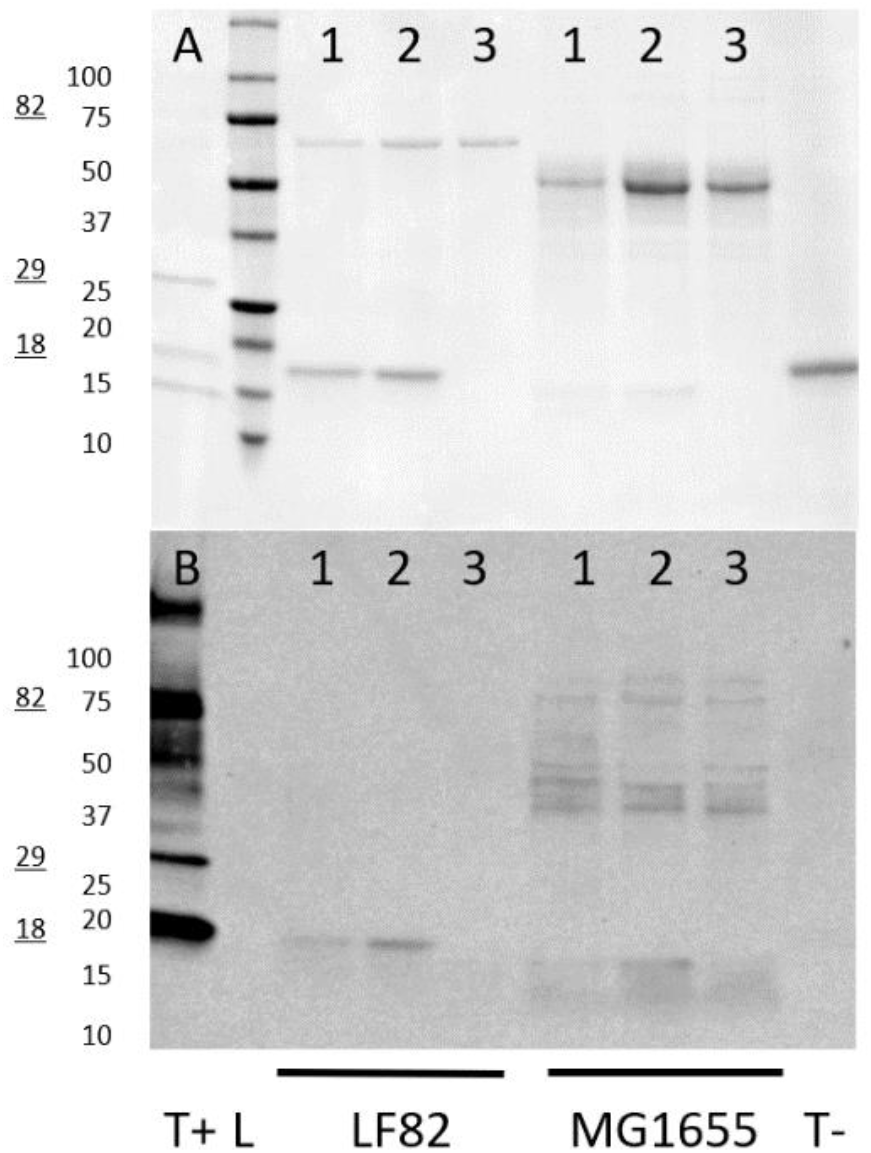
Biotin-hydrazide-labelling of glycoprotein sugars. Glycoproteins on PVDF blots were biotin-hydrazide labelled and detected by streptavidin-HRP and enhanced chemiluminescence. (A) Ponceau S staining; (B) Streptavidin-HRP visualization. Glycoprotein ladder is used as a positive control (T+, 18 kDa, 29 kDa, and 82 kDa glycoprotein bands), while soybean trypsin inhibitor is the negative control (T-). Samples loaded for each strain are: 1 – acid hydrolyzed protein extract, 2 – guanidine HCl-treated surface extract, 3 – untreated surface extract. LF82 – AIEC LF82, MG1655 – *E. coli* K-12 MG1655.

**Figure 12.**
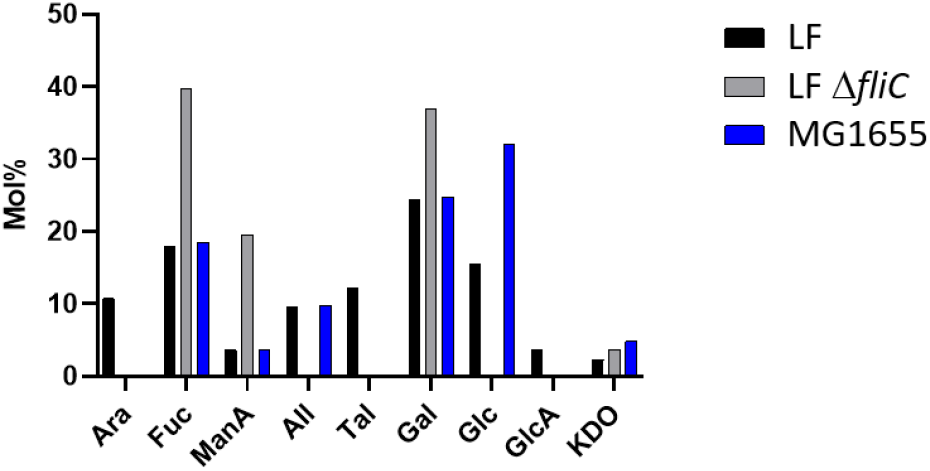
Compositional analysis of flagella/pili-enriched surface protein extracts. LF – AIEC LF82, LF Δ*fliC* – LF82 Δ*fliC*, MG1655 – *E. coli* K-12 MG1655. **Ara** – arabinose, **Fuc** – fucose, **ManA** – mannuronic acid, **All** – allose, **Tal** – talose, Gal – galactose, Glc – glucose, **GlcA** – glucuronic acid, KDO - ketodeoxyoctonic acid. Highlighted sugars are those not found in O83 O-antigen.

Further, fimbria-enriched surface protein extracts of AIEC LF82 and *E. coli* K-12 MG1655 underwent compositional analysis via acid hydrolysis and gas chromatography-mass spectrometric analysis (GC-MS) (Fig. 12). The results indicate the presence of monosaccharides present in the LPS inner and outer core of both strains (including glucose - Glc, galactose – Gal, N-acetyl-glucosamine - GlcNAc, and ketodeoxyoctonic acid - KDO), however, the analysis demonstrated the presence of non-LPS sugars as well, including allose (All), talose (Tal), mannuronic acid (ManA), glucuronic acid (GlcA), arabinose (Ara), and fucose (Fuc). Thus, while we and others may have initially thought the samples to be contaminated with LPS, the presence of non-LPS sugars implies the presence of alternative glycans / glycoproteins. Further, comparison of the wild-type and the Δ*fliC* mutant extracts, certain sugars are no longer detected in the absence of FliC, namely arabinose, allose, talose, glucose, and glucuronic acid. Alternatively, the FliC-mutant was associated with increases in fucose, mannuronic acid, and galactose.

As GuHCl treatment results in the most efficient depolymerization of FimA oligomers, having the highest detection by interaction with ConA, this approach was employed to prepare FimA for binding affinity determination by thermal shift. The interaction of a dilution series of FimA with a fixed concentration of NHS-RED-tagged ConA led to the determination of the K_D_ of interaction of approximately 40μM.

### Enzymatic treatments

#### PNGase F or mannosidase

Based on the previous staining by lectins recognizing mannose, more information was sought on the identity or nature of the glycan linkage. Enzymatic treatments using glycosidases were implemented in an attempt to cleave the attached glycan. Both PNGase F and mannosidase treatments did not result in a demonstrable change in ConA staining of the FimA subunit (Fig. 13A).

**Figure 13.**
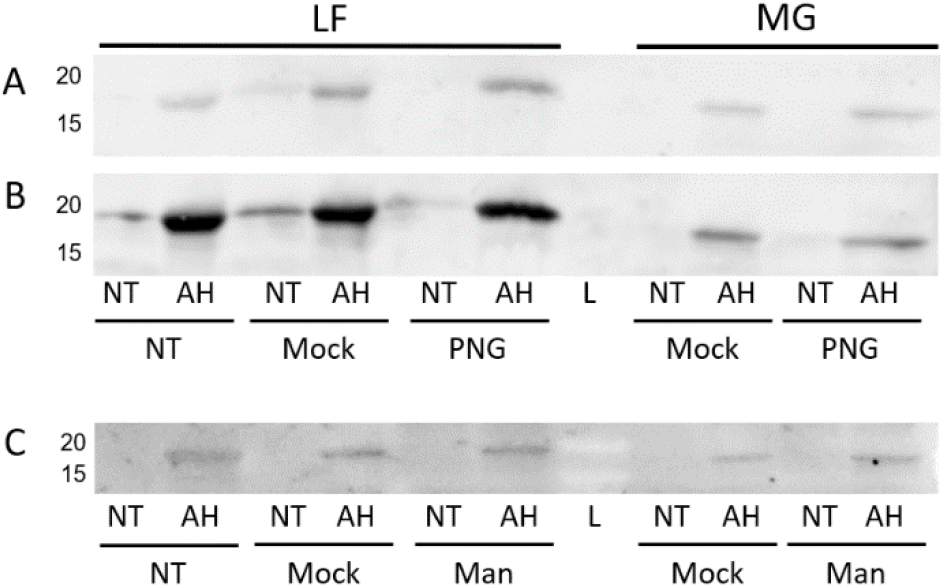
ConA lectin probing following PNGase or mannosidase treatment of released *E. coli* surface proteins. LF: AIEC LF82, MG: *E. coli* MG1655. NT: Non-acid hydrolyzed, AH: Acid hydrolyzed, L: Molecular weight ladder, Mock: Mock-treated sample, Man: Mannosidase-treated sample. (A) PNGase F treatment probed with ConA. (B) Anti-type I fimbria staining of membrane (A). (C) Mannosidase treatment probed with ConA.

However, interestingly, incubation with either of these enzymes resulted in a significant increase in the detection of type I fimbria by Western blot in comparison to the mock-treated samples (Fig. 14). The exact mechanism has not yet been identified. While treatment with these enzymes did not manage to diminish the ConA binding, it has demonstrated that the treatments have an effect on the behavior of the fimbrial polymers, by increasing the amount of fimbrial polymer to enter into the SDS-PAGE gel. In samples treated with PNGase F, novel bands were detected by means of the stain-free technology in the Bio-Rad TGX SDS-PAGE gels (Fig. 15). However, while bands were excised and analyzed by proteomics, the polymeric nature of FimA/pili or flagella/FliC resulted in the primary result in each band being FimA or FliC, while neither protein is typically detected by the stain-free system owing to their lack of tryptophan residues. Notably, following PNGase F treatment, GNA staining was significantly reduced, nearly eliminating the smear and was associated with the appearance of a GNA-stained band of approximately 31 kDa (Fig. 10A, lanes 2 and 6), however, the identity of this protein has not yet been determined, nor has a PNGase F-released glycan been identified by standard glycomics analyses.

**Figure 14.**
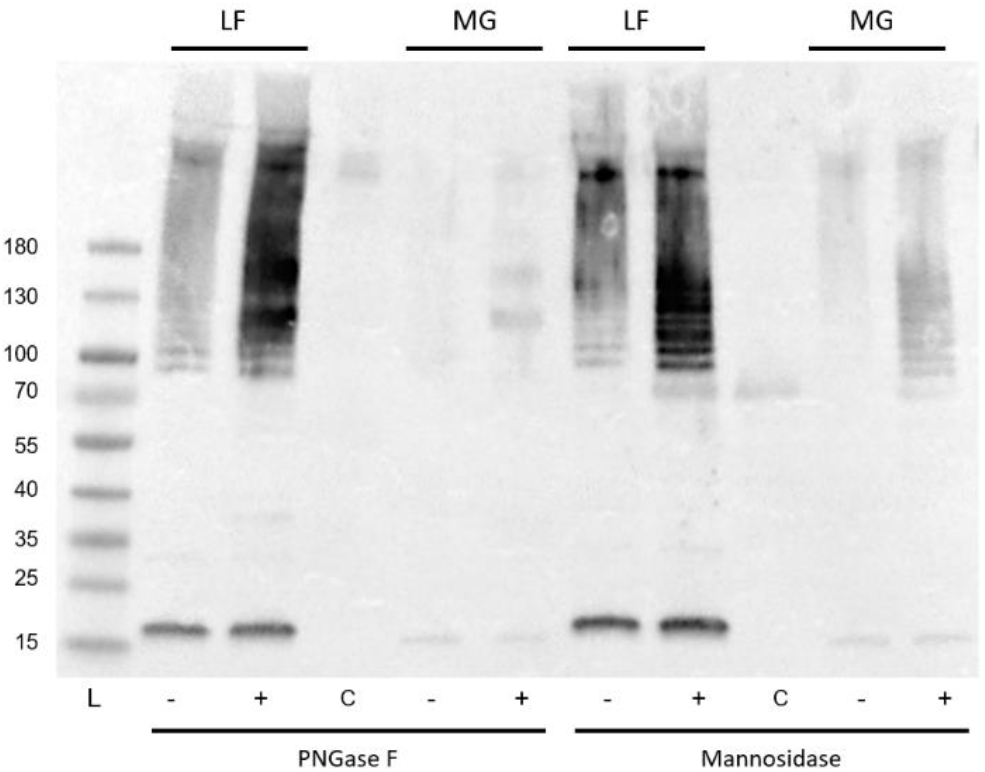
Anti-type I fimbria staining following PNGase F or mannosidase treatment of released *E. coli* surface proteins. PNGase F or mannosidase treatment of released *E. coli* surface proteins increased anti-type I pili detection. L: PageRule protein ladder; LF: AIEC LF82; MG: *E. coli* MG1655; - denotes mock-treated, + denotes enzyme treated, C denotes enzyme-only control.

**Figure 15.**
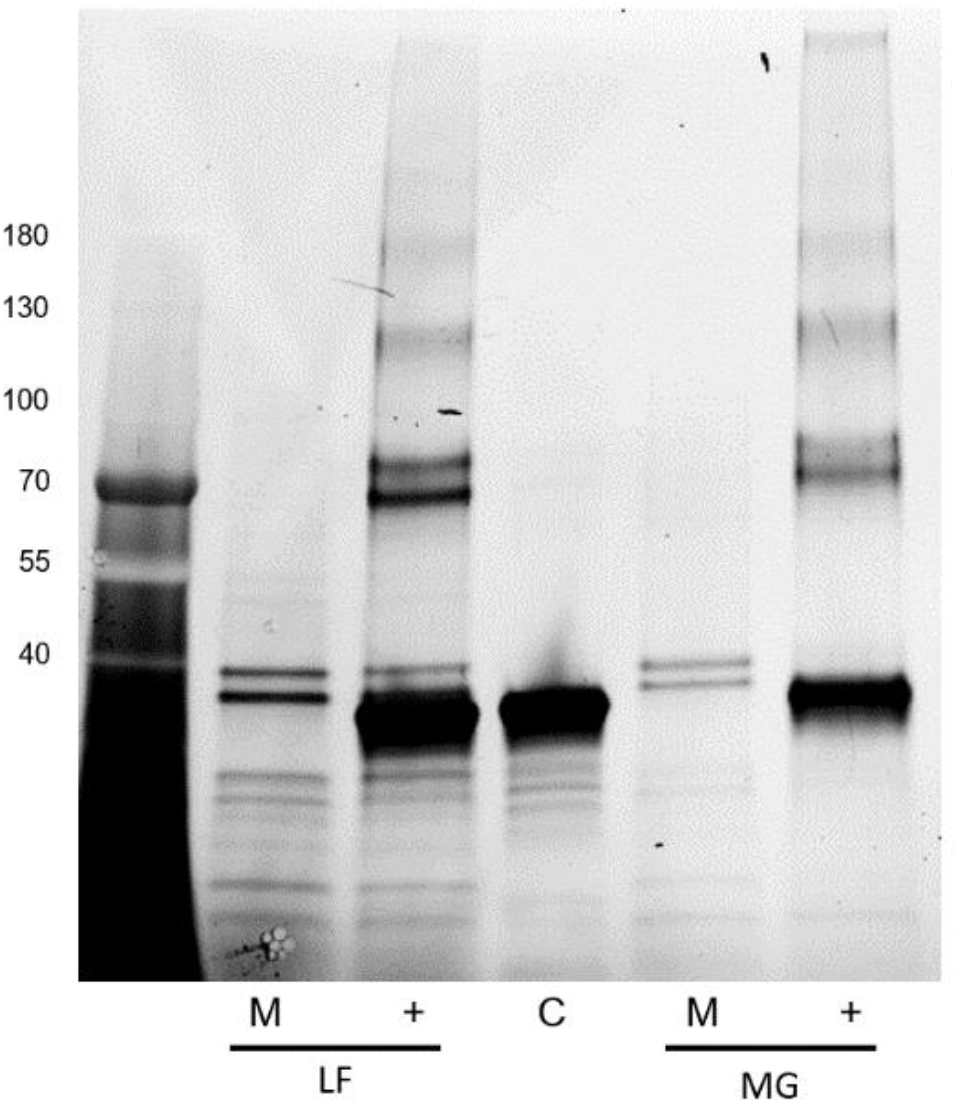
Stain-free detection following PNGase F treatment of released *E. coli* surface proteins. Far left: Molecular weight ladder. LF: AIEC LF82, MG: *E. coli* MG1655, M denotes mock-treated sample, + denotes PNGase F-treated sample, and C denotes the enzyme only control.

While PNGase F treatment of samples results in a generally altered behavior of type I fimbriae, Western blot of identically treated samples with anti-FliC antibody results in a novel ∼32 kDa band, implying a susceptibility of the FliC protein to degradation by PNGase F (Fig. 16).

**Figure 16.**
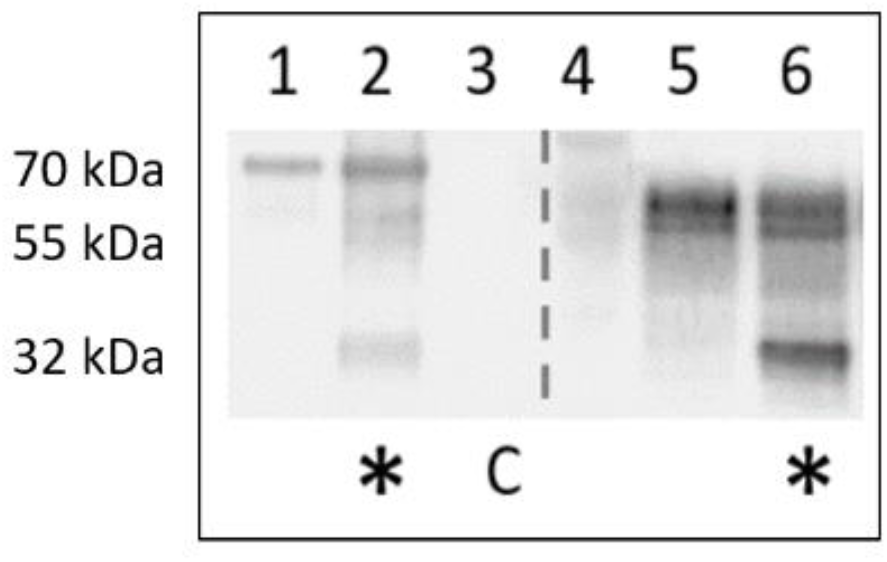
Anti-FliC detection in PNGase F-treated surface protein extracts of AIEC LF82 and *E. coli* K-12 MG1655. Lanes 1,5 – mock treatment with denaturation buffer, enzyme replaced by H2O. Lanes 2, 6 – PNGase-F-treated ^(*^). C denotes PNGase F enzyme-only control.

### Chemical treatments

Alkaline treatment was next tested on samples of surface proteins of LF82 for its ability to release *O*-linked glycans. Following overnight incubation in alkaline conditions (pH >10) at 45°C, samples were assayed by lectin blot with ConA to determine if the treatment would reduce the intensity of the binding. However, the opposite finding was observed, and following alkaline incubation staining with ConA of FimA subunits was actually increased (not shown) likely indicating a more efficient release of FimA monomers for detection. Furthermore, while it was considered that the FimA monomers would be robust enough to survive beta-elimination, treatment results in complete degradation of the FimA protein, hindering the determination of whether the glycan is *O*-linked by SDS-PAGE mobility shift or lectin staining.

## Discussion

To our knowledge, this represents the first study demonstrating the presence of a glycan on the FimA subunit of *Escherichia coli* type I fimbriae via Western blot. Previous studies have indicated the presence of reducing sugars^3^ and speculated on their identities^11^ in preparations of purified pilin, but have not gone so far as to demonstrate the interaction of a specific lectin with FimA subunits, and to explore means by which to remove the associated glycan.

Type I fimbriae, along with flagella, are surface proteins extending into the external environment (approximately 1 μm and 5-10 μm, respectively)^28,29^ much further than that of LPS (3-5 nm or 17-37 nm, depending on the presence of O-antigen)^30^ and their expression can be modulated by contact with host structures^31^. As such, fimbriae represent an initial point of contact of the bacterium with many structures, including other surrounding bacteria, the intestinal epithelium, the overlying mucus barrier, as well as host immune cells^1^. Unfortunately, the study and characterization of type I fimbriae is inherently difficult.

Firstly, type I fimbriae are polymeric structures, which impedes the resolution of the involved subunits. In fimbrial preparations, the minor subunits, FimF and FimG, are reported to be present at a ratio of <1:100 to the major subunit, FimA, with quantities of FimH even more scarce^32^. As such, their study is limited by either the requirement for large batch cultures (often >5L)^26^, or their heterologous expression^33^, which risks artificial or non-natural conditions. Secondly, the unique property of the fimbrial FimA subunit assembly hinders detachment and purification of the major subunit. In turn, FimA subunit release often requires acid hydrolysis, further hindering the study of the ensemble of subunits at once. To overcome this obstacle, studies have relied on deletion mutants or the overexpression of specific subunits^33^. As studies have focused on the role of individual subunits, the behavior and composition of intact polymers has been largely overlooked, despite early interest primarily around the 1970s^8,9,34^.

In the current study we identified a common fimbrial behavior in which intact fimbrial polymers migrate in gradient SDS-PAGE gels. This was detected both in the presence and absence of the high molecular weight smear of fimbrial polymer, as a well-defined band, pointing to a minimal unit size of FimA oligomer of approximately 90 kDa. This structure, likely representing a pentameric assembly, resists SDS, beta-mercaptoethanol, and heat treatment, but can be further depolymerized with harsher treatments to release FimA subunits. As the fimbrial subunits FimF, FimG, and FimH readily dissociate from the structure following SDS treatment at elevated temperatures^5^, it could be proposed that each or a combination of FimF, FimG, or FimH are located along the pilus^8,34^ and once released by SDS result in a minimal FimA homopolymer size of approximately 90 kDa, present as a common minimal size of type I fimbrial structure maintained across several laboratory and pathobiont strains studied here. Further evidence of smaller, interspersed spans of FimA is put forth by Eshdat *et al*., in which they determine that their GuHCl fimbrial preparations contain the lectin component, FimH, at 1%. For a 28 kDa protein, this extrapolates to 1 FimH subunit in 2800 kDa, or 1 FimH subunit per ∼165 FimA subunits, given a size of 17 kDa and ignoring the presence of the other two subunits, FimF and FimG^9^. At present, there is mounting evidence of the distribution of FimF, FimG, and FimH along the fimbrial shaft, yet most models incorporate these subunits solely at the fimbrial tip.

Earlier papers studying type I fimbriae are often focused on fimbrial subunits, and accordingly the SDS-PAGE gels are restricted or cropped to <100 kDa in most cases^5^, and as such the presence of the fimbrial smear and the size at which it stops is not discussed^32^. Type I fimbria purification typically includes 3 rounds of MgCl_2_ precipitation, aiding in the removal of fimbrial fragments, as well as other contaminants. This possibly explains the absence of the FimA-pentamer in most papers investigating type I fimbria, as it may bias the purification to the larger, intact structures. Of note, we observed that the amount of time permitted for re-solubilization of MgCl_2_ precipitated fimbria could influence the presence of the high molecular weight type I fimbria polymer smear (higher degree of solubility), or favor the presence of the 90 kDa FimA oligomer, as well as FimH presence in the pellet that required an extended period to resolubilize (results unpublished). Further of note is that Abraham *et al*. produced monoclonal antibodies, recognizing either a specific FimA polymeric quaternary structure, but not the dissociated FimA subunits, or solely the dissociated FimA subunits, but not the fully assembled structure. This indicates both, the recognition of larger structures (repeating intervals, in a spiral motif), or hidden antigens, only detected upon dissociation of fimbriae^10^. In the current study, the anti-type I fimbria serum was produced in a rabbit host, lacking the specificity of monoclonal antibodies, but allowing the detection of various forms of FimA-bearing fimbrial constructs and the individual FimA subunits.

Following the observation of ConA staining of released FimA subunits, but not of the non-hydrolyzed type I fimbria samples, it was hypothesized that the glycan may be inaccessible due to either steric hindrance or internal localization in the tight helical fimbrial structure. This is in agreement with the findings of Abraham *et al*., in which glycerol treatment to uncoil fimbriae or guanidine hydrochloride treatment to induce dissociation of type I fimbriae permitted antibody recognition of previously hidden epitopes^10,35^. Accordingly, we hypothesized given that acid hydrolysis and release of FimA subunits from the polymeric fimbriae is required to expose the attached glycan to enable lectin binding, acid hydrolysis should be included before enzymatic treatment, with the aim to remove the newly exposed glycan. However, pre-treatment by acid hydrolysis before PNGase F enzymatic treatment failed to decrease ConA recognition of the FimA subunit, implying either i) the enzymatic treatments were not successful despite the exposure of the FimA glycan, or ii) re-neutralization of samples prior to treatment results in re-folding of the FimA subunit and protection of the associated glycan, although acid hydrolysis is generally considered irreversible.

The exact role of the glycans linked to FimA has not yet been elucidated, but based on chemical and enzymatic treatments to remove sugars unexpectedly increasing the quantity of fimbriae migrating into SDS-PAGE gels, it is hypothesized that the currently unidentified sugars may play a role in *E. coli* type I pilus and flagellar stability. Glycans are typically vulnerable to acid hydrolysis, yet McMichael and Ou demonstrated that the detection of reducing sugars on type I fimbriae requires acid hydrolysis^3^. Additionally, their study implied the presence of a non-*O*-linked glycan present on fimbriae, possibly representing an alternative glycan moiety, proposed to be an *N*-glycan, based on their alkaline hydrolysis results. However, our results following digestion with PNGase F or mannosidase indicate that the FimA-associated glycan is either not an *N*-glycan, or a non-classical *N*-glycan resisting PNGase F cleavage. Further regarding PNGase F activity, we have demonstrated the susceptibility of FliC to be degraded following incubation with PNGase F. This results in a band of approximately 32 kDa staining with anti-FliC antibodies. At present it has not been determined if it is a degradation product, or has been released by disruption of an unidentified interaction. Sequence analysis of the FliC protein demonstrates the potential presence of an *N*-glycan attachment site based on the bacterial *N*-glycosylation sequon D/E-X-N-X-S/T (X ≠ Pro), where Asn (N) is the attachment site^36^. Cleavage of the protein at this site would produce an appropriately sized fragment, but this remains to be confirmed, as PNGase F is not commonly associated with protein cleavage.

Despite lectin staining of surface protein extracts enriched for fimbriae and flagella indicating the presence of mannose or glucose, treatment with a mannosidase did not reduce the lectin staining of ConA (GNA staining confounded by staining of mannosidase enzyme), nor did lectin inhibition with alpha-methylmannose or glucose (100mM). Interestingly, PNGase F or mannosidase treatment increased the staining of type I fimbriae in treated samples compared with mock-treated controls.

To rule out the potential for aberrant behavior of the lectins, a methodology which directly conjugated biotin to an oxidized glycan was employed. The results indicate the distinct staining of the FimA subunits in the LF82 samples, but not FliC. Conversely, the *E. coli* K-12 MG1655 sample demonstrated staining for FimA subunits, as well as separate bands varying between 30-80 kDa. Secondly, compositional analysis revealed the presence of mannuronic acid in the enriched fimbria sample of LF82, which was augmented in the flagella-deficient mutant. Given the potential for acidic saccharides, Alcian blue staining was employed, first staining positive for dot blots for pili-enriched samples, and later on total protein blots across several strains, revealing conserved staining of a ∼36 kDa band, as well as strain specific staining. This staining implies the presence of acidic- or muco-polysaccharides associated with surface proteins in the panel of strains screened. The exact identities and function require further research.

As *E. coli* do not possess a characteristically typical glycosylation pathway, the origin of the glycan is hypothesized to come from a non-specialized synthetic pathway, potentially associated with the assembly of LPS. Thorsing *et al*.^24^ have investigated the presence of heptose on the YghJ protein secreted by *E. coli* H10407, and in doing so have inactivated the *hldE* gene in MG1655, used as the protein-expression strain. The previous works of Boysen *et al*^37^ demonstrated that the pathogenic strain was more likely to glycosylate surface proteins of the H10407 strain. However, it draws attention to the fact that it was deemed necessary to express the YghJ protein in the MG1655 Δ*hldE* strain, inferring the ability of the common *E. coli* K-12 MG1655 strain to modify proteins with heptose.

Isogenic mutants for the genes involved in the LPS synthesis pathway are often associated with an absence or reduction in the expression of surface organelles, including flagella and fimbriae. More specifically, this involves the genes required for the synthesis or addition of the LPS inner core saccharides. While knockout of those genes involved in the attachment of 3-deoxy-D-manno-oct-2-ulosonic acids (Kdo) to lipid A are lethal, those involved in the attachment of the subsequent heptoses produce a phenotype referred to as ‘deep rough’ linked to colony appearance^38,39^. The first gene involved in heptose addition, *hldE* (formerly *waaE* or *rfaE*) encodes the HepI transferase^40^. *HldE* mutants in ETEC H10407 are non-flagellated and display altered piliation^41^. Recently, HldE has demonstrated the capacity to glycosylate non-LPS structures, including YghJ, which is a secreted metalloprotease able to degrade the intestinal mucin layer, thus facilitating colonization. As LPS synthesis and expression and presence of fimbria and flagella are closely interconnected, the investigation of the potential activity of LPS enzymes in their modification via isogenic mutants is hindered.

The limitations of our study lie in the fact that classic glycomics and glycoproteomics analyses have thus far failed to identify a fimbrial or FimA-associated glycan. However, this may be explained, at least partly, following the enzymatic and chemical treatments in the current study, and that FimA is inherently resistant to trypsin digestion^35^. As it was not possible to directly confirm the presence of a fimbria-associated glycan(s), indirect biochemical means were employed. Of note, based on the findings of Abraham *et al*.^35^, sample preparation for glycoproteomics may benefit from the inclusion of glycerol, in order to augment the fimbriae subunits susceptibility to trypsin digestion, however, necessitating later removal of the glycerol from the digested peptides.

Additionally, bacterial surface protein glycan modifications are often in lower abundance than those of eukaryotic cells, and require enrichment steps for their detection. As an example, in the study by Boysen *et al*.^37^ in which glycopeptides were enriched, the YghJ protein was originally found to possess 4 amino acid residues with *O*-linked heptose, while in a subsequent study it was found to be hyperheptosylated bearing 54 occupied sites. Notably, despite the previous efforts of groups to identify *E. coli* glycoproteins^14^, the use of lectins specific to either more standard *N*-glycans or *O*-glycans will have missed those proteins modified with heptose.

Despite the exact identity of the FimA subunit modification remaining currently unknown, the study has cast light on the varying behavior of the FimA subunits, while they are typically regarded as a homopolymer of identical subunits. However, the current study demonstrates the presence of a pentameric FimA structure, which when treated with guanidine hydrochloride results in a FimA tetramer which demonstrates varying susceptibility reducing agents. Accordingly, this sequence of treatments highlights the differences in resistance of FimA oligomers, as well as novel means to selectively dissociate FimA oligomers to distinct sizes.

In summary, the results of the current study demonstrate the presence of posttranslational modification of type I fimbriae and highlight further areas of research interest regarding the surface proteins of *E. coli*. Future research will focus on determining the exact identity of the FimA-associated glycan, as well as its importance to bacterial survival, colonization, or infection.

## Materials and methods

Magnesium chloride, dithiothreitol (DTT), iodoacetamide (IAA), biotinylated wheat germ agglutinin (WGA) lectin, FITC-conjugated antirabbit IgG antibody, and biotin- and HRP-conjugated concanavalin A (ConA) lectin were obtained from Merck Sigma (France). HRP-conjugated galanthus nivalis lectin (GNA) was obtained from USBiological (CliniSciences, Nanterre; G1044-25). HRP-conjugated anti-rabbit antibody was obtained from Cell Signaling Technology via Ozyme (France). Primary rabbit anti-FliC antibody (ab93713) was obtained from AbCam, and the anti-type I fimbria serum was a kind gift from Dr. Annie Brée (Université de Tours, France)

### Bacterial strains and growth conditions

Ampicillin-erythromycin–resistant *E. coli* strain LF82, isolated from a chronic ileal lesion of a CD patient^42^, was used as the AIEC reference strain in the current study. The AIEC strain LF82 and the laboratory strain *E. coli* K-12 MG1655 were routinely cultured 48 hours in Luria-Bertani (LB) broth at 37°C under static conditions to promote fimbriae expression. Additional *E. coli* K-12, deletion mutants, and AIEC strains used in the screening experiment are part of the M2iSH laboratory strain collection and are found in Table I.

**Table I.**
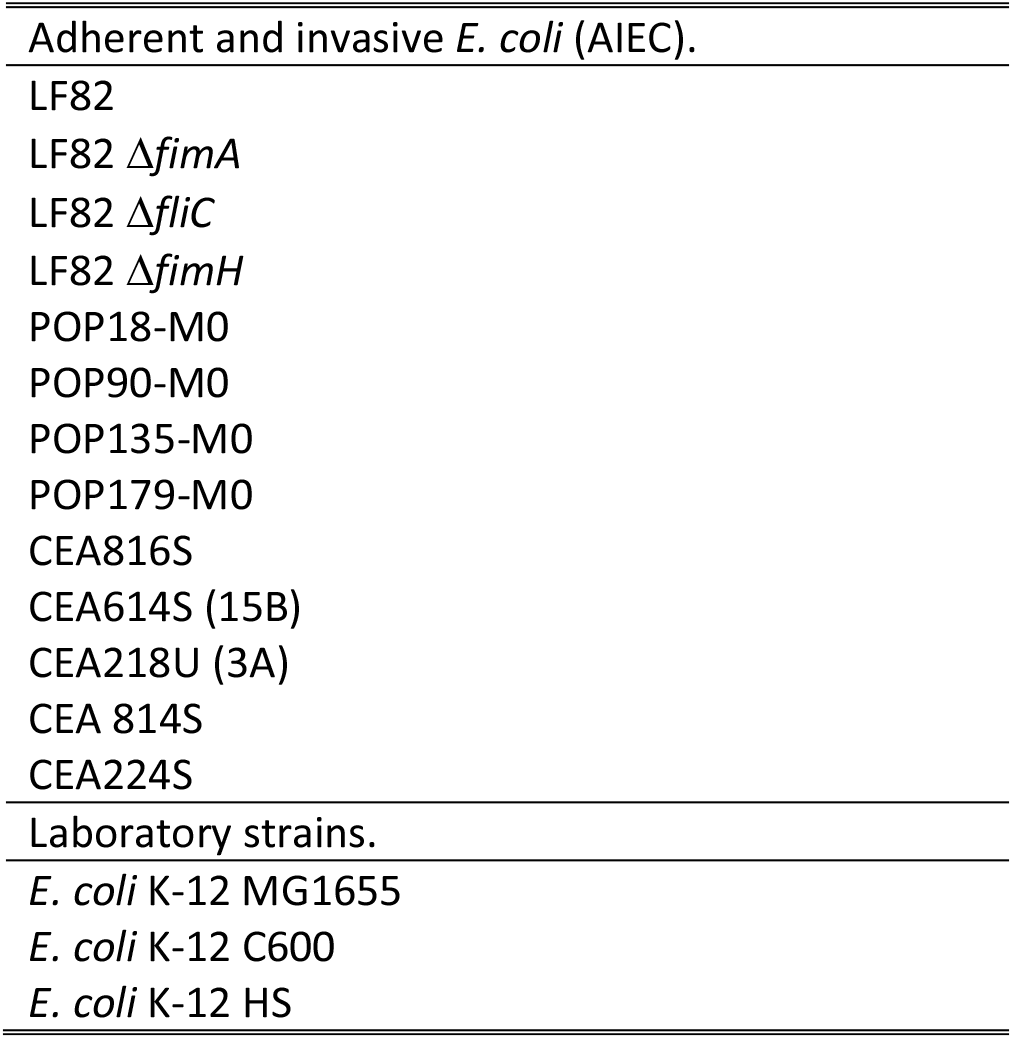
Bacterial strains used in the current study.

### Protein purification

Bacterial cultures were grown under static conditions for 48 hours at 37°C to promote fimbriae expression. Following growth, cultures were incubated at 4°C for 30 min, and then harvested by centrifugation at 10,000 x g, 4°C x 10 min. Bacterial pellets were then washed with cold PBS, and finally resuspended at a final volume of 45ml PBS / 500ml culture. Bacterial surface proteins were sheared by mechanical force using an Ultraturrax blender with 5 pulses at 16000 rpm of 1 min on ice, with 1 min rest periods between pulses. Intact bacteria and debris were then pelleted by centrifugation at 10 000 x g, for 15 min at 4°C, and the supernatant containing released proteins conserved. Alternatively, to limit or avoid potential fragmentation of released flagella and pili, extraction of surface proteins was implemented as per Barnich et al, (2003). Briefly, instead of mechanical shearing, collected and washed bacteria were resuspended in PBS and heated at 60°C for 20 min with vigorous shaking. This extraction method was found to yield equivalent amounts of flagella and type I fimbriae relative to mechanical shearing, while enabling higher throughput of samples.

When global analysis of released proteins was required, released surface proteins were precipitated with trichloroacetic acid (TCA; 10% final concentration). Following 30 min at 4°C, samples were collected by centrifugation at 15,000 x g for 15 min, washed twice in 100% ice cold acetone, and resuspended in 1/1000 volume milli-Q water and stored at 4°C until analyzed, as freeze-thaw has been demonstrated to fragment pili^8^. Relative enrichment of flagella and type I fimbriae was carried out by precipitation in 30% ammonium sulphate, with overnight incubation under gentle agitation at 4°C, and collection by centrifugation as above. More strict purification of pili and flagella was achieved by 2 rounds of precipitation in 100 mM MgCl_2_ ≥ 2 h at 4°C, recovery of precipitated proteins by centrifugation, and resuspension in milli-Q water.

### SDS-PAGE and Western blotting

*E. coli* surface protein extracts were characterized by SDS-PAGE. 4-20% stain-free TGX gradient gels were used to allow the detection of both non-acid-hydrolyzed and acid hydrolyzed samples simultaneously. For analysis, 9 μl of surface extract was mixed with 4x Laemmli buffer, and heated at 99°C for 5 min prior to loading. Acid hydrolysis of samples was performed by the addition of 0.2 μl of 3.7% HCl to the 12 μl sample, followed by heating at 99°C for 5 min. Samples were then neutralized with ∼0.6μl 1M NaOH, and heated at 99°C for 5 min prior to loading. Stain-free protein detection was performed according to Bio-Rad instructions. Western blot analysis of samples was performed with the TurboBlot system, transferring the SDS-PAGE gel to PVDF using the high molecular weight program. Following transfer, blots were dried under a stream of warm air and then re-activated in methanol prior to blocking and detection. Blots were blocked in protein-free TBS buffer (Thermo Scientific) for ≥ 1 h. To visualize FimA or FliC, blots were incubated with either antibodies raised against *E. coli* type I fimbria or FliC (diluted 1:10 000 in protein-free tris-buffered saline (TBS) blocking solution (Fisher Scientific, France)) for 1 h at room temperature, respectively. Blots were then washed 3x in TBS+0.1% Tween 20 (TBST), and incubated 1 h with anti-rabbit-HRP or anti-rabbit-IgG-FITC (1:10,000; Sigma, France) in protein-free TBS blocking buffer. Blots were washed 4 × 10 min in TBST, following by direct visualization of FITC, or enhanced chemiluminescence via Clarity (Bio-rad) in Bio-rad MP chemidoc. Lectin staining was performed on membranes blocked as for antibody detection.

Galanthus nivalis lectin (GNA), Concanavalin A (ConA) or wheat germ agglutinin lectin (WGA) conjugated with horseradish peroxidase (HRP) were diluted to 10 μg/ml in protein-free TBS and incubated for ≥ 1 h at room temperature. Blots were washed 4 × 10 min with TBST and then visualized by enhanced chemiluminescence using the Clarity Max (Bio-Rad) substrate.

Alcian blue staining was performed following protein transfer to nitrocellulose membranes. Membranes were washed with deionized water, then incubated in 1% (w/v) Alcian blue in 3% (v/v) acetic acid, pH 2.5 for 30 min. Membranes were then extensively washed in deionized water to remove unbound dye.

### *Enzymatic and chemical treatment of* E. coli surface proteins

To further probe the nature of the attached glycans, protein aliquots were subjected to PNGase F or a broad range mannosidase (α1,2/3/6) (New England Biolabs, France) digestion according to the manufacturer’s specifications, with the exception that incubations were carried out overnight for 18hrs. Mock treatments were identical, maintaining the initial denaturation step, but substituting water for enzyme. Following incubation, if necessary, samples were concentrated by means of chloroform:methanol precipitation and resuspended in 12 μl H_2_O for SDS-PAGE analysis. Where acid hydrolysis was performed before and after enzymatic treatments, with the aim of facilitating enzyme access to potential glycans, chloroform:methanol precipitation was used to precipitate samples and adjust sample volumes. To do so, samples were precipitated, acid hydrolyzed, neutralized, precipitated, and resuspended in the original volume of H_2_O.

Alkaline treatment was carried out, as per McMichael and Ou^3^, on samples to investigate whether the attached glycan(s) were susceptible to alkaline hydrolysis. Surface protein extracts were adjusted to pH > 10 and incubated 16 h at 45°C. Control samples remained neutral, but were treated 16 h at 45°C.

Further, B-elimination was carried out similarly to the alkaline treatment, however, including 1 M sodium borohydride as reductant in the 100 mM NaOH, pH 12 solution. This treatment is typically very harsh on proteins, but pili and FimA subunits display unusually highly resistant properties. The resulting solution was neutralized with 5% acetic acid dropwise until the fizzing stopped. The solution was then dialyzed against water (MWCO 3,000 Da) overnight with 3 changes in order to remove salts.

For reduction and alkylation, samples were incubated in 10mM dithiothreitol at 56°C for 30 min, followed by incubation with 30 mM iodoacetamide at room temperature for 30 min. Samples were then precipitated by the chloroform:methanol method to remove the chemical reagents and resuspended in the original volume.

Guanidine hydrochloride treatment was performed on chloroform:methanol precipitated samples resuspended in saturated GuHCl (∼8.6 M) in water for 2 h at 37°C. The guanidine solution is first heated to >37°C to allow saturation, as a saturated solution at 20°C corresponds to ∼6 M, rather than 8.6 M at 37°C. Samples were dialyzed extensively against water to remove GuHCl and then concentrated prior to further use.

Following enzymatic or chemical treatments, samples were analyzed by antibody and lectin blotting, to determine the impact on the presence/detection of pili, FimA subunits, or glycan staining.

### Nanotemper Monolith X interaction analysis of FimA monomers and ConA

Following optimization of FimA monomer release, the FimA-ConA interaction was characterized by spectral shift using capillary tubes. PBS, pH 6.8 was used as reaction buffer. FimA was prepared at 10 μM, and the NHS-red-tag kit (Nanotemper, Germany) was used to label ConA (Sigma, France), which was diluted according to manufacturer’s instructions.

### Hydrazine-biotin tagging of PVDF immobilized proteins

PVDF membranes obtained using the above methodology were washed in milli-Q water followed by glycan oxidation via incubation in 10 mM sodium metaperiodate in 100 mM sodium acetate buffer, pH 5.5 for 20 min in the dark at 4°C with gentle agitation. Membranes were washed extensively with water to remove residual periodate, followed by incubation with 5 mM biotin-hydrazide (Sigma, France) for 60 minutes, at ambient temperature with agitation and protected from light. Membranes were blocked with 3% non-fat milk for 1.5 h followed by probing with streptavidin-HRP (1:1000) for 1 h. Labelled glycans were visualized with Clarity ECL (Bio-Rad) and imaged with a Bio-Rad MP Chemidoc. A glycoprotein ladder (Candy Cane ladder; Invitrogen) served as positive control, and soybean trypsin inhibitor, as well as other molecular weight marker proteins (Bio-Rad precision plus blue) as negative controls.

### Monosaccharide compositional analysis

Compositional analysis of monosaccharides was conducted on surface protein preparations enriched for type I fimbriae and flagella. Lyophilized extracts of surface proteins underwent methanolysis in 500 μL of 0.5 M HCl in anhydrous methanol at 80°C for 16 hours. Methanolic HCl was evaporated and the resulting products were per-acetylated in 200 μL of acetic anhydride and 50 μL of pyridine overnight at room temperature. Reagents were evaporated and the sample was dissolved in chloroform prior to GC-MS analysis.

## Acknowledgments

This study was made possible through the generous financing of the Université Clermont Auvergne encouraging the mobility of international researchers. This work was supported by the “Ministère de l’Enseignement Supérieur, de la Recherche et de l’Innovation (MESRI), Inserm (Institut national de la santé et de la recherche médicale; UMR1071), INRAE (Institut national de recherche en agriculture, alimentation et environnement; USC 1382), the Agence Nationale de la Recherche of the French government through the program “Investissements d’Avenir” (IDEX-ISITE initiative 16-IDEX-0001 CAP 20-25 project of the University of Clermont Auvergne), the ANR (French National Research Agency), and the National Program “Microbiote” Inserm.

We would like to thank the kind gift of anti-FimF-C and anti-FimG-C antibodies from Prof. David Thanassi of Stony Brook University, New York, USA. We would also like to acknowledge the extensive advice and conversations with Stephanie Archer-Hartmann and Dr. Parastoo Azadi of the Complex Carbohydrate Research Centre, Georgia, USA, as well as Dr. Dimitrios Latousakis of the Quadram Institute, Norwich, UK, and Dr. Priscilla Branchu of INRAE, Toulouse, France.

## References

1. Avalos Vizcarra I, Hosseini V, Kollmannsberger P, et al. How type 1 fimbriae help Escherichia coli to evade extracellular antibiotics. Sci Rep. 2016;6(1):18109. doi:10.1038/srep18109

2. Kaper JB, Nataro JP, Mobley HLT. Pathogenic Escherichia coli. Nat Rev Microbiol. 2004;2(2):123–140. doi:10.1038/nrmicro818

3. McMichael JC, Ou JT. Structure of common pili from Escherichia coli. J Bacteriol. 1979;138(3):969–975.

4. Hanson MS, Hempel J C C Brinton J. Purification of the Escherichia coli type 1 pilin and minor pilus proteins and partial characterization of the adhesin protein. J Bacteriol. Published online August 1988. doi:10.1128/jb.170.8.3350-3358.1988

5. Hanson MS, Brinton CC. Identification and characterization of E. coli type-1 pilus tip adhesion protein. Nature. 1988;332(6161):265–268. doi:10.1038/332265a0

6. Brinton CC. The structure, function, synthesis and genetic control of bacterial pili and a molecular model for DNA and RNA transport in Gram negative bacteria. Trans N Y Acad Sci. 1965;27(8 Series II):1003–1054. doi:10.1111/j.2164-0947.1965.tb02342.x

7. Krogfelt KA, Bergmans H, Klemm P. Direct evidence that the FimH protein is the mannose-specific adhesin of Escherichia coli type 1 fimbriae. Infect Immun. 1990;58(6):1995–1998. doi:10.1128/iai.58.6.1995-1998.1990

8. Ponniah S, Endres RO, Hasty DL, Abraham SN. Fragmentation of Escherichia coli type 1 fimbriae exposes cryptic D-mannose-binding sites. J Bacteriol. 1991;173(13):4195–4202.

9. Eshdat Y, Silverblatt FJ, Sharon N. Dissociation and reassembly of Escherichia coli type 1 pili. J Bacteriol. 1981;148(1):308–314. doi:10.1128/jb.148.1.308-314.1981

10. Abraham SN, Hasty DL, Simpson WA, Beachey EH. Antiadhesive properties of a quaternary structure-specific hybridoma antibody against type 1 fimbriae of Escherichia coli. J Exp Med. 1983;158(4):1114–1128. doi:10.1084/jem.158.4.1114

11. Palacios Pelaez R, Martinez Garate A, Martinez Quesada J. Type 1 and type p fimbriae-adhesins isolated from novel e. coli strains, process for their preparation and uses thereof. Published online October 29, 2002. https://patents.google.com/patent/US6471966B1/en

12. Duncan MJ, Mann EL, Cohen MS, Ofek I, Sharon N, Abraham SN. The Distinct Binding Specificities Exhibited by Enterobacterial Type 1 Fimbriae Are Determined by Their Fimbrial Shafts*. J Biol Chem. 2005;280(45):37707–37716. doi:10.1074/jbc.M501249200

13. Dreux N, Denizot J, Martinez-Medina M, et al. Point mutations in FimH adhesin of Crohn’s disease-associated adherent-invasive Escherichia coli enhance intestinal inflammatory response. PLoS Pathog. 2013;9(1):e1003141. doi:10.1371/journal.ppat.1003141

14. Wang Z xiu, Deng R ping, Jiang HW, et al. Global identification of prokaryotic glycoproteins based on an Escherichia coli proteome microarray. Chatterji D, ed. PLoS ONE. 2012;7(11):e49080. doi:10.1371/journal.pone.0049080

15. Neuhaus A, Selvaraj M, Salzer R, et al. Cryoelectron microscopy reveals two distinct type IV pili assembled by the same bacterium. Nat Commun. 2020;11(1):2231. doi:10.1038/s41467-020-15650-w

16. Benz I, Schmidt MA. Glycosylation with heptose residues mediated by the aah gene product is essential for adherence of the AIDA-I adhesin. Mol Microbiol. 2001;40(6):1403–1413. doi:10.1046/j.1365-2958.2001.02487.x

17. Logan SM, Hui JPM, Vinogradov E, et al. Identification of novel carbohydrate modifications on Campylobacter jejuni 11168 flagellin using metabolomics-based approaches: Flagellin glycosylation in C. jejuni. FEBS J. 2009;276(4):1014–1023. doi:10.1111/j.1742-4658.2008.06840.x

18. Grass S, Lichti CF, Townsend RR, Gross J, St. Geme JW. The Haemophilus influenzae HMW1C Protein Is a Glycosyltransferase That Transfers Hexose Residues to Asparagine Sites in the HMW1 Adhesin. Stebbins CE, ed. PLoS Pathog. 2010;6(5):e1000919. doi:10.1371/journal.ppat.1000919

19. Schirm M, Soo EC, Aubry AJ, Austin J, Thibault P, Logan SM. Structural, genetic and functional characterization of the flagellin glycosylation process in Helicobacter pylori: Glycosylation of Helicobacter flagellin. Mol Microbiol. 2003;48(6):1579–1592. doi:10.1046/j.1365-2958.2003.03527.x

20. Horzempa J, Held TK, Cross AS, et al. Immunization with a Pseudomonas aeruginosa 1244 Pilin Provides O-Antigen-Specific Protection. Clin Vaccine Immunol. 2008;15(4):590–597. doi:10.1128/CVI.00476-07

21. Hartley MD, Morrison MJ, Aas FE, Børud B, Koomey M, Imperiali B. Biochemical Characterization of the O-Linked Glycosylation Pathway in Neisseria gonorrhoeae Responsible for Biosynthesis of Protein Glycans Containing N, N ′-Diacetylbacillosamine. Biochemistry. 2011;50(22):4936–4948. doi:10.1021/bi2003372

22. Hopf PS, Ford RS, Zebian N, et al. Protein Glycosylation in Helicobacter pylori: Beyond the Flagellins? Neyrolles O, ed. PLoS ONE. 2011;6(9):e25722. doi:10.1371/journal.pone.0025722

23. Lu Q, Li S, Shao F. Sweet Talk: Protein Glycosylation in Bacterial Interaction With the Host. Trends Microbiol. 2015;23(10):630–641. doi:10.1016/j.tim.2015.07.003

24. Thorsing M, Krogh TJ, Vitved L, et al. Linking inherent O-Linked Protein Glycosylation of YghJ to Increased Antigen Potential. Front Cell Infect Microbiol. 2021;11. Accessed March 15, 2022. https://www.frontiersin.org/article/10.3389/fcimb.2021.705468

25. Barnich N, Boudeau J, Claret L, Darfeuille-Michaud A. Regulatory and functional cooperation of flagella and type 1 pili in adhesive and invasive abilities of AIEC strain LF82 isolated from a patient with Crohn’s disease: Flagella and adherent-invasive E. coli. Mol Microbiol. 2003;48(3):781–794. doi:10.1046/j.1365-2958.2003.03468.x

26. Jones CH, Pinkner JS, Roth R, et al. FimH adhesin of type 1 pili is assembled into a fibrillar tip structure in the Enterobacteriaceae. Proc Natl Acad Sci U S A. 1995;92(6):2081–2085.

27. Jaipuri FA, Collet BYM, Pohl NL. Synthesis and Quantitative Evaluation of Glycero-D-manno-heptose Binding to Concanavalin A by Fluorous-Tag Assistance. Angew Chem Int Ed. 2008;47(9):1707–1710. doi:10.1002/anie.200704262

28. Namba K, Yamashita I, Vonderviszt F. Structure of the core and central channel of bacterial flagella. Nature. 1989;342(6250):648–654. doi:10.1038/342648a0

29. Miller E, Garcia T, Hultgren S, Oberhauser AF. The Mechanical Properties of E. coli Type 1 Pili Measured by Atomic Force Microscopy Techniques. Biophys J. 2006;91(10):3848–3856. doi:10.1529/biophysj.106.088989

30. Strauss J, Burnham NA, Camesano TA. Atomic force microscopy study of the role of LPS O-antigen on adhesion of E. coli: AFM STUDY OF E. coli LPS. J Mol Recognit. 2009;22(5):347–355. doi:10.1002/jmr.955

31. Sheikh A, Rashu R, Begum YA, et al. Highly conserved type 1 pili promote enterotoxigenic E. coli pathogen-host interactions. Yang R, ed. PLoS Negl Trop Dis. 2017;11(5):e0005586. doi:10.1371/journal.pntd.0005586

32. Krogfelt KA, Klemm P. Investigation of minor components of Escherichia coli type 1 fimbriae: protein chemical and immunological aspects. Microb Pathog. 1988;4(3):231–238. doi:10.1016/0882-4010(88)90073-3

33. Alonso-Caballero A, Schönfelder J, Poly S, et al. Mechanical architecture and folding of E. coli type 1 pilus domains. Nat Commun. 2018;9(1):2758. doi:10.1038/s41467-018-05107-6

34. Abraham SN, Goguen JD, Sun D, Klemm P, Beachey EH. Identification of two ancillary subunits of Escherichia coli type 1 fimbriae by using antibodies against synthetic oligopeptides of fim gene products. J Bacteriol. 1987;169(12):5530–5536. doi:10.1128/jb.169.12.5530-5536.1987

35. Abraham SN, Land M, Ponniah S, Endres R, Hasty DL, Babu JP. Glycerol-induced unraveling of the tight helical conformation of Escherichia coli type 1 fimbriae. J Bacteriol. 1992;174(15):5145–5148. doi:10.1128/jb.174.15.5145-5148.1992

36. Kowarik M, Young NM, Numao S, et al. Definition of the bacterial N-glycosylation site consensus sequence. EMBO J. 2006;25(9):1957–1966. doi:10.1038/sj.emboj.7601087

37. Boysen A, Palmisano G, Krogh TJ, Duggin IG, Larsen MR, Møller-Jensen J. A novel mass spectrometric strategy “BEMAP” reveals Extensive O-linked protein glycosylation in Enterotoxigenic Escherichia coli. Sci Rep. 2016;6(1):32016. doi:10.1038/srep32016

38. Raetz CRH, Whitfield C. Lipopolysaccharide Endotoxins. Annu Rev Biochem. 2002;71(1):635–700. doi:10.1146/annurev.biochem.71.110601.135414

39. Nakao R, Ramstedt M, Wai SN, Uhlin BE. Enhanced Biofilm Formation by Escherichia coli LPS Mutants Defective in Hep Biosynthesis. Dobrindt U, ed. PLoS ONE. 2012;7(12):e51241. doi:10.1371/journal.pone.0051241

40. Desroy N, Moreau F, Briet S, et al. Towards Gram-negative antivirulence drugs: New inhibitors of HldE kinase. Bioorg Med Chem. 2009;17(3):1276–1289. doi:10.1016/j.bmc.2008.12.021

41. Maigaard Hermansen GM, Boysen A, Krogh TJ, Nawrocki A, Jelsbak L, Møller-Jensen J. HldE Is Important for Virulence Phenotypes in Enterotoxigenic Escherichia coli. Front Cell Infect Microbiol. 2018;8:253. doi:10.3389/fcimb.2018.00253

42. Darfeuille-Michaud A, Neut C, Barnich N, et al. Presence of adherent Escherichia coli strains in ileal mucosa of patients with Crohn’s disease. Gastroenterology. 1998;115(6):1405–1413. doi:10.1016/s0016-5085(98)70019-8

